# *Crk/Crkl* regulates early angiogenesis in mouse embryos by accelerating endothelial cell maturation

**DOI:** 10.1101/2023.07.12.548782

**Authors:** Lijie Shi, Hansoo Song, Bin Zhou, Bernice E. Morrow

**Author notes:** Corresponding author: Bernice E. Morrow, Email address Lijie Shi, Email address Address: 1301 Morris Park Avenue, Price Building room 475, Department of Genetics, Albert Einstein College of Medicine, Bronx, NY 10461, USA.

## Abstract

**Rationale:** Ubiquitously expressed cytoplasmic adaptors CRK and CRKL mediate multiple signaling pathways in mammalian embryogenesis. They are also associated with cardiovascular defects occurring in Miller-Dieker syndrome and 22q11.2 deletion syndrome, respectively. The embryonic mesoderm contributes to the formation of the cardiovascular system, yet the roles that *Crk* and *Crkl* play there are not understood on a single cell level.

**Objectives:** To determine functions of *Crk* and *Crkl* in the embryonic mesoderm during early mouse vascular development. Secondly, we will examine the molecular mechanisms responsible for early embryonic endothelial cell (EC) defects by performing single cell RNA-sequencing (scRNA-seq) and *in vivo* validation experiments.

**Methods and Results:** Inactivation of both *Crk* and *Crkl* together using *Mesp1^Cre^* resulted embryonic lethality with severe vascular defects. Although vasculogenesis appeared normal, angiogenesis was disrupted both in the yolk sac and embryo proper, leading to disorganized vascular networks. We performed scRNA-seq of the *Mesp1^Cre^* mesodermal lineage and found that there was upregulation of a great number of angiogenesis and cell migration related genes in ECs in the mutants, including NOTCH signaling genes such as *Dll4* and *Hey1*. Further bioinformatic analysis of EC subpopulations identified a relative increase in the number of more differentiated angiogenic ECs and decrease in EC progenitors. Consistent with this, we identified an expansion of *Dll4* expressing cells within abnormal arteries, *in vivo*. Also, our bioinformatic data indicates that there is dysregulated expression of lineage genes that promote EC differentiation causing accelerated cell fate progression during EC differentiation.

**Conclusions:** Our results show that *Crk* and *Crkl* are crucial for regulating early embryonic angiogenesis. Combined inactivation of *Crk/Crkl* caused precocious EC maturation with an increase of atypical differentiated angiogenic ECs and failed vascular remodeling. This is in part due to increased NOTCH signaling and altered expression of cell migration genes.

## Introduction

Ubiquitously expressed cytoplasmic adaptor proteins, CRK and CRKL (CRK like), mediate many essential signaling pathways in response to extracellular stimuli.^1–3^ In humans, *CRK* and *CRKL* are closely related genes and are located on chromosomes 17p13.3 and 22q11.2, respectively. Haploinsufficiency of genes in these regions are associated with congenital malformation disorders termed, Miller-Dieker syndrome (MDS) ^4–6^ and 22q11.2 deletion syndrome (22q11.2DS) ^7,8^, respectively. Some affected individuals with MDS ^4,5,9–13^ and most with 22q11.2DS ^14,15^ have cardiovascular defects. It is possible that *CRK* and *CRKL* contribute to congenital cardiovascular disease in these individuals. This is supported by studies in mouse models demonstrating that global inactivation of either *Crk* or *Crkl* results in mid-gestational embryonic lethality due to various aspects of vascular and intracardiac defects.^16–20^ Mouse model studies have also shown that *Crk* and *Crkl*, have shared functions in several biological contexts including embryonic development.^16,20–23^ This report focuses on the functions of *Crk* and *Crkl* in the mesoderm on vascular development in early mouse embryogenesis.

Mesodermal cells contribute to the vasculature of the yolk sac and embryo including the formation of endothelial cells (ECs), vascular smooth muscle cells (SMCs) as well as pericytes.^24–26^ The ECs form the inner lining of all blood vessels, of which most are surrounded by SMCs and pericytes. During early embryogenesis, formation of blood vessels in the yolk sac and embryo occurs by vasculogenesis and angiogenesis processes. Vasculogenesis involves *de novo* blood vessel formation from EC progenitors, which mainly occurs before the heart starts beating (mouse embryonic day, E8.0-8.25, 3-4 somites); while angiogenesis occurs after vasculogenesis and is a process by which blood vessels expand by creation of new branches followed by coalescence and remodeling to form a functional vascular network.^27–30^ One of the first steps of angiogenesis involves specification of ECs to different cell types.

The formation of endothelial tip and stalk cells among ECs is a sign of the beginning of spouting angiogenesis.^31^ Tip cells are located at the leading end of vascular sprouts and are characterized by their position as well as the presence of long filopodia, high migratory ability and lower proliferative rate; while stalk cells follow behind the tip cells, and in comparison to tip cells, have fewer filopodia, highly proliferative rate and establish tight junctions between cells.^31^ Dysregulation of signaling pathways or genes crucial for regulating specification of tip and stalk cells or their functions will cause defective angiogenesis and/or vascular remodeling.^31–39^ Multiple signaling pathways have been studied for their crucial roles in regulating vasculogenesis and angiogenesis processes, such as VEGF, FGF, TGFβ signaling in vasculogenesis, and VEGF and NOTCH signaling in angiogenesis.^40^

The embryonic mesoderm forms at gastrulation and transient expression of the *Mesp1* in these mesodermal cells marks a significant part of the future cardiovascular system including ECs.^24,25,41^ Loss of *Crk* and *Crkl* in the mesoderm in the mouse using *Mesp1^Cre^*has been reported to cause heart looping and yolk sac vascular remodeling defects, however mice died too early to further investigate their functions.^16^ Additionally, detailed morphological studies have not yet been performed to understand the basis for yolk sac vascular defects. It also remains unknown whether the embryos themselves have vascular defects. Finally, the cellular and molecular mechanisms that explain vascular phenotypes on a single cell basis need to be explored. Therefore, this study is focused upon the role of ECs in the early embryonic mesoderm during vascular development that require the shared functions of *Crk* and *Crkl*.

To achieve this, we used the *Mesp1^Cre^* allele to inactivate both floxed alleles of *Crk* and *Crkl*. We found grossly normal vasculogenesis both in the mutant yolk sac and embryo, but severe disorganization of blood vessels comprising the vascular network. To investigate molecular mechanisms, we performed single cell RNA sequencing (scRNA-seq) of the embryonic *Mesp1* lineage at E8.5 and focused on ECs. We found up-regulation of multiple angiogenesis as well as cell migration related genes including NOTCH signaling genes *Dll4* in ECs in the mutants. Further we found loss of *Crk/Crkl* caused accelerated and atypical EC maturation with increased *Dll4* expressing angiogenic cells and potentially defective vessel remodeling in the mutants. Additionally, our data shows that dysregulation of *Mesp1* lineage genes that are known to promote EC differentiation may contribute to precocious EC maturation. In summary, our data provide potential molecular mechanisms for how *Crk/Crkl* regulate EC differentiation, angiogenesis and vascular remodeling in the early mammalian embryo.

## Methods

### Mouse strains

The *Mesp1^Cre/+^*,^24^ *Rosa26 ^GFP/+^* (GFP *^f/+^*),^42^ *Crk ^f/f^*,^18^ *Crkl ^f/f^* ^20^ lines used in this study were previously described. The *Mesp1^Cre/+^* and *Rosa26 ^GFP/+^* mice were maintained in the Swiss Webster background. The *Crk ^f/f^*, *Crkl ^f/f^* and *Crk ^f/f^;Crkl ^f/f^* mice were intercrossed and maintained in a mixed genetic background (C57BL/6, 129SvEv and Swiss Webster) for approximately 3-4 years. All mouse studies were performed according to the ethical guidelines (Protocol #00001034) approved by the Institutional Animal Care of Use Committee at Albert Einstein College of Medicine (https://www.einsteinmed.edu/administration/animal-care-use-committee/).

### Whole mount immunofluorescence staining (WHIF) of embryos and yolk sac

The embryos along with their yolk sac at desired stages (E8.5: 3-10 somites; E9.5: 19-24 somites) were isolated in phosphate-buffered saline (PBS; Corning #21-030-CV), fixed using 4% paraformaldehyde (PFA) overnight at 4°C on a rocker, then washed twice in PBS at 4°C to remove the PFA. The embryos (and/or yolk sac) were then dehydrated through a methanol series (25%, 50%, 75%, 100% x 2) on ice and stored at -20°C. Embryos (and/or yolk sac) used for WHIF were either from those after PFA fixation and PBS wash, or those from the freezer which were rehydrated through a methanol series (75%, 50%, 25%, PBS x 2) on ice.

All WHIF steps were performed at 4°C on a rocker. Embryos were washed with 0.1% Triton X-100 in PBS (PBST) overnight, then blocked using 5% donkey serum (Sigma #D9663; diluted in 0.1% PBST) overnight. After washing with 0.1% PBST (5 times in daytime and one time overnight), embryos were incubated with primary antibodies for two days. The embryos were then washed with 0.1% PBST and then were incubated with secondary antibodies together with DAPI for two days. Next, embryos were washed by 0.1% PBST, dehydrated through a methanol series (25%, 50%, 75%, 100% x 2), and then cleared by Benzyl Alcohol and Benzyl Benzoate (Benzyl Alcohol: Benzyl Benzoate = 1:2 in volume) before imaging. Embryos were imaged using a Nikon CSU-W1 spinning disk confocal microscope from Einstein Analytical Imaging Facility (https://www.einsteinmed.edu/research/shared-facilities/analytical-imaging-facility/). Three-dimensional images of embryos were visualized and analyzed using ImarisViewer 9.6.0 software.

Primary antibodies used in this study included Rat anti-PECAM1 (BD Biosciences #550274), Goat anti-GFP (Abcam #ab6673), Rabbit anti-pH3 (Sigma #631257). Secondary antibodies used included Donkey anti-Goat IgG(H+L) Alexa Fluor™ 488 (Invitrogen #A11055), Donkey anti-Rat IgG(H+L) AF555 (Southern Biotech #643032) and Donkey anti-Rabbit IgG(H+L) Alexa Fluor™ 647 (Invitrogen #A31573).

TUNEL assays were performed after secondary antibody staining by using the *In Situ* Cell Death Detection Kit TMR red (Millipore #12156792910). Embryos were incubated for 4 hours at room temperature (RT) on rocker to develop signals for cells undergoing apoptosis. Notch intracellular domain (NICD) staining was performed on paraffin sections using methods previously described.^20^ Tyramide signal amplification was performed for NICD staining.

### Whole mount RNAscope in situ hybridization of embryos

Whole mount RNAscope assays were performed in 96-well plates using Multiplex Fluorescent Reagent kit v2 (Advanced Cell Diagnostics #323100). Probes used for RNAscope were from Advanced Cell Diagnostics including *Acta2* (#319531), *Nkx2-5* (#428241), *Meox1* (#530641-C2), *Tagln* (#480331-C2), *Pecam1* (#316721-C3), *Tbx1* (#481911), *Tbx5* (#519581-C2), *Isl1* (#451931-C2), *Dll4* (#319971-C2), *Etv2* (#447481).

Embryos were processed in the same way as those used for WHIF according to manufacturer’ instructions with modifications of the length of incubations depending on embryonic stages. All wash steps for RNAscope assays were performed at RT for 3 minutes (3 times) on a rocker, and all incubation steps in an oven were performed at 40°C. Embryos were washed with 0.1% Tween20 in PBS (0.1% PBS-T), permeabilized by using Protease plus (Advanced Cell Diagnostics #322331) for 15 minutes, then washed by 0.01% PBS-T. Next, embryos were incubated with prewarmed probes (volume ratio for probes C1:C2:C3=50:1:1) in oven for overnight, then washed by 0.01% Tween20 in 0.2 x Saline Sodium Citrate (0.01% T-0.2xSSC). Then embryos were fixed in 4% PFA for 10 minutes, incubated with Amp1, Amp2, Amp3 for 30, 30, 15 minutes at 40°C, respectively, with a wash by 0.01% T-0.2xSSC between each step. Tyramide Signal Amplification (TSA) steps were performed for detecting fluorescent signals. TSA solutions included TSA-Fluorescein (Akoya Biosciences #NEL741001KT), TSA-CY3 (Akoya Biosciences #SAT704A001EA) and TSA-CY5 (Akoya Biosciences #SAT705A001EA). To reveal C1 probes, embryos were incubated with HRP-C1 at 40°C for 15 minutes, washed with 0.01% T-0.2xSSC, incubated with chosen TSA solution at 40°C for 30 minutes for developing signals, washed with 0.01% T-0.2xSSC, then incubated with HRP-Blocker at 40°C for 15 minutes to stop the amplification reaction. Signals for C2 and C3 probes were revealed following the previous steps by using HRP-C2, HRP-C3 for C2, C3 probes, respectively. After washing with 0.01% T-0.2xSSC, embryos were then stained with DAPI overnight at 4°C on a rocker to visualize nuclei. Next, embryos were dehydrated through a methanol series gradient, cleared with Benzyl Alcohol and Benzyl Benzoate and then were ready for imaging. Images were taken using a Nikon CSU-W1 spinning disk confocal microscope and analyzed by ImarisViewer 9.6.0 software as described above.

### Single cell RNA-seq (scRNA-seq) data generation

Embryos at E8.5 (3-8 somite) were collected in pre-chilled cold PBS and then stored in a 96-well plate with Dulbecco’s Modified Eagle Medium (DMEM; GIBCO #11885084) on ice. Genotyping was performed immediately to identify the genotypes of embryos, which took approximately 2 hours. Next, embryos with desired genotypes were selected and the rostral part of embryos including the hearts were micro-dissected in ice-cold PBS. Then the micro-dissected tissues with same genotype were pooled in the same 1.5 milliliter (ml) Eppendorf tube and incubated with 0.7 ml of 0.25% Trypsin-EDTA (Gibco #25200056) consisting of 50 Unit/ml DNase I (Millipore #260913-10MU) at RT for 7 minutes. Next, 78 microliters (μl) of heat inactivated Fetal Bovine Serum (FBS; ATCC #30-2021) was added to stop the reaction, and gentle pipetting was performed to help dissociate tissues to single cells. The dissociated cells were then centrifuged at 300 x gravity (g) for 5 minutes at 4°C and the supernatants were discarded. Next, cells were resuspended in 0.6 ml PBS (without calcium & magnesium; Corning #21-031-CV) containing 10% FBS on ice and filtered through a 100 μm cell strainer. One microliter of 1mM DAPI (Thermo Fisher Scientific #D3571) was added before FACS, and GFP positive plus DAPI negative cells were collected in 0.5 ml PBS (without calcium & magnesium)/10% FBS using a BD FACSAria II system for FACS in the Einstein Flow Cytometry Core Facility (https://www.einsteinmed.edu/research/shared-facilities/cores/10/flow-cytometry/). Next, sorted cells were centrifuged at 300 x g for 5 minutes at 4°C and resuspended in 50 μl of PBS (without calcium & magnesium) containing 10% FBS.

Sorted cells were then loaded in the 10x Chromium instrument (10x Genomics) using the dual index Chromium Next GEM Single Cell 3’ v3.1 kit for library preparation. Next, DNA libraries were sequenced using an Illumina NovaSeq system (Genewiz from Azenta Life Sciences) with paired-end, 150 bp read length. Two biological replicates of control (*Mesp1 ^Cre/+^;Crk ^f/+^;Crkl ^f/+^*) and cKO (*Mesp1 ^Cre/+^;Crk ^f/f^;Crkl ^f/f^*) samples were prepared. Embryos for Control 1 (Ctrl1) and cKO1 were collected from same isolated litters, and embryos for Ctrl2 and cKO2 were collected from another several isolated litters.

### Single cell RNA-seq data analysis

Alignment of scRNA-seq reads to the mouse reference genome assembly, mm10-2020-A, and generation of gene-by-cell count matrices were performed using Cell Ranger v6.0.1 software (10x Genomics). The Seurat v4.1.1 package ^43^ was used to filter and cluster the individual scRNA-seq data per sample. The filtering parameters for cells and clusters were chosen according to the recommendations from the RISC v1.0 package.^44^ This software package was used to integrate all scRNA-seq datasets including two replicates of control and cKO samples from the Seurat individual objects. All cells from Seurat filtering were included in RISC analysis.^44^ Expression data was normalized for the integrated data and cells were re-clustered with control #2 as the reference based on parameters recommended by RISC software. Differentially expressed genes (DEGs) between control and cKO embryos were determined using RISC software for single or combined cell clusters with an adjusted P-value cut off of <0.05 and |log_e_^(fold change)^| cut off of 0.25. Heatmaps of DEGs were prepared using RISC software. Gene ontology (GO) analysis of the DEGs was performed using the ToppFun application within ToppGene Suite (https://toppgene.cchmc.org/enrichment.jsp).^45^ Protein-Protein interactions were analyzed by using STRING v11.5.

For the cell trajectory analysis, Cell Ranger output files were processed using Velocyto v0.17.17 software^46^ to generate loom files. These loom files and individual Seurat UMAPs were imported to be used by scVelo v0.2.3^47^ software to map cell fates based on RNA velocities from a dynamical model. CellRank v1.5.1^48^ was used to calculate absorption probabilities, which are the probabilities of each individual cell to progress to a certain fate. Potential driver genes were identified by CellRank based on dynamical behavior and likelihood to fit within the dynamical model. Velocities and absorption probabilities were plotted using Matplotlib v3.3.4 and Scanpy v1.7.1. Heatmaps of driver genes were plotted using seaborn v0.11.2 software. The original Seurat cluster identities of the cells were retained and used to track the cell clusters where they originated from when analyzing RISC UMAP plots.

### Flow cytometry

Cells were dissociated from the yolk sac of desired genotypes at E8.5 using the method and reagents the same as for cells prepared for scRNA-seq. After adding 10% FBS to stop reaction, gentle pipetting was performed. Then cells were gently passed through 28G1/2 needle twice to help dissociate tissues to single cell suspensions which were subject to following antibody staining. PE-conjugated anti-PECAM1 (CD31; Invitrogen #12-0311-82) and APC-conjugated anti-cKit (CD117; Invitrogen #17-1171-82) antibodies were used. All staining and washing were performed in the 2%FBS/PBS either on ice or on rocker at 4°C. DAPI (Invitrogen) was added before FACS to label dead cells. Flow cytometry data were collected by the same machine used for collecting scRNA-seq cells.

### Statistics

The proportion of cells in each cluster per sample was calculated from the integrated data. The two-proportion Z-test was calculated in RStudio and was used to evaluate the statistical significance of the change in cell proportions between control and cKO samples with a P-value cut off at 0.05. The two-proportion Z-test was also used for evaluating the statistical significance of the change in the number of up- and down-regulated DEGs per cell cluster.

### Data availability

The scRNA-seq datasets for this study are available in the NCBI Gene Expression Omnibus (GEO) database with accession number of GSE220583.

## Results

### Loss of Crk and Crkl in the Mesp1 lineage results in abnormal vascular formation in the yolk sac and embryo proper

To investigate the functions of *Crk* and *Crkl* in the mesoderm in early mouse development, we conditionally inactivated one or both alleles of *Crk* and *Crkl* using the *Mesp1^Cre^* allele.^18,20,24^ For phenotypic analysis, we termed the conditional mutants with inactivation of one allele of *Crk* and one allele of *Crkl* as 2-allele cKO embryos, one allele of *Crk* and two alleles of *Crkl* (and vice versa) as 3-allele cKO embryos, and finally, inactivation of both alleles of *Crk* and *Crkl* as 4-allele cKO embryos.

We found that the 2-allele cKO mice were viable, healthy and fertile. In contrast, most of 3- and 4-allele cKO embryos died between E10.5 and E11.5 (Table S1). At E8.5, we did not observe any obvious differences among control (Cre negative) and 2-, 3- and 4-allele cKO mutants regarding the structure of the yolk sac capillary plexus (Fig. S1; minimal N=5 for each genotype). At E9.5 and E10.5, we observed a poorly remodeled yolk sac capillary plexus in 3- and 4-allele cKO mutants versus normal remodeled large vessels in both control and 2-allele cKO embryos (Fig. S1). This indicates that there was failure of yolk sac vascular remodeling in 3- and 4-allele cKO embryos. The vascular findings in the yolk sac were observed by whole mount light microscopy previously in 3-allele cKO embryos.^16^ Our 3- and 4-allele cKO embryos were similar in size to each other, but smaller in size compared to either 2-allele or control embryos at E9.5 and E10.5 (Fig. S1). It was previously reported that 4-allele cKO embryos had severe growth arrest after gastrulation preventing analysis of the vasculature.^16^ However our 4-allele cKO embryos died at E11.5, which allowed us to study the function of both *Crk/Crkl* in the early embryonic angiogenesis. The differences of the timing of embryonic lethality of 4-allele cKO embryos in our study may be due to differences in the genetic background of our mice that were used.

To visualize the vasculature network better in the yolk sac, we performed whole mount immunofluorescence staining using an anti-PECAM1 antibody on the control (*Mesp1^Cre/+^;GFP ^f/+^*) versus 3- and 4-allele cKO embryos at E8.5 (Fig. 1A-B) and E9.5 (Fig. 1C-D; Fig. 1, minimal N=3 for each genotype). PECAM1 is a widely used marker for ECs. At E8.5, we observed a capillary plexus network both in the yolk sac of control and 4-allele cKO embryos (Fig. 1A-B); however, it seems that there were less dense capillary plexuses in 4-allele cKO embryos (Fig. 1B) suggesting abnormal vascular development. At E9.5, we found that the yolk sac of control embryos exhibited well delineated large blood vessels (Fig. 1C), while the 4-allele cKO embryos had only immature vascular plexuses without the presence of large vessels (Fig. 1D). To investigate further whether there were vascular abnormalities at E8.5, we quantified PECAM1 positive cells by flow cytometry and observed a reduction of PECAM1 positive cells in the yolk sac of 3- and 4-allele cKO embryos compared to the controls (Fig. S2; Fig. 1E; Cre negative for control; minimal N=3 for each genotype), indicating that there were slightly less ECs contributing to the yolk sac.

**Figure 1.**
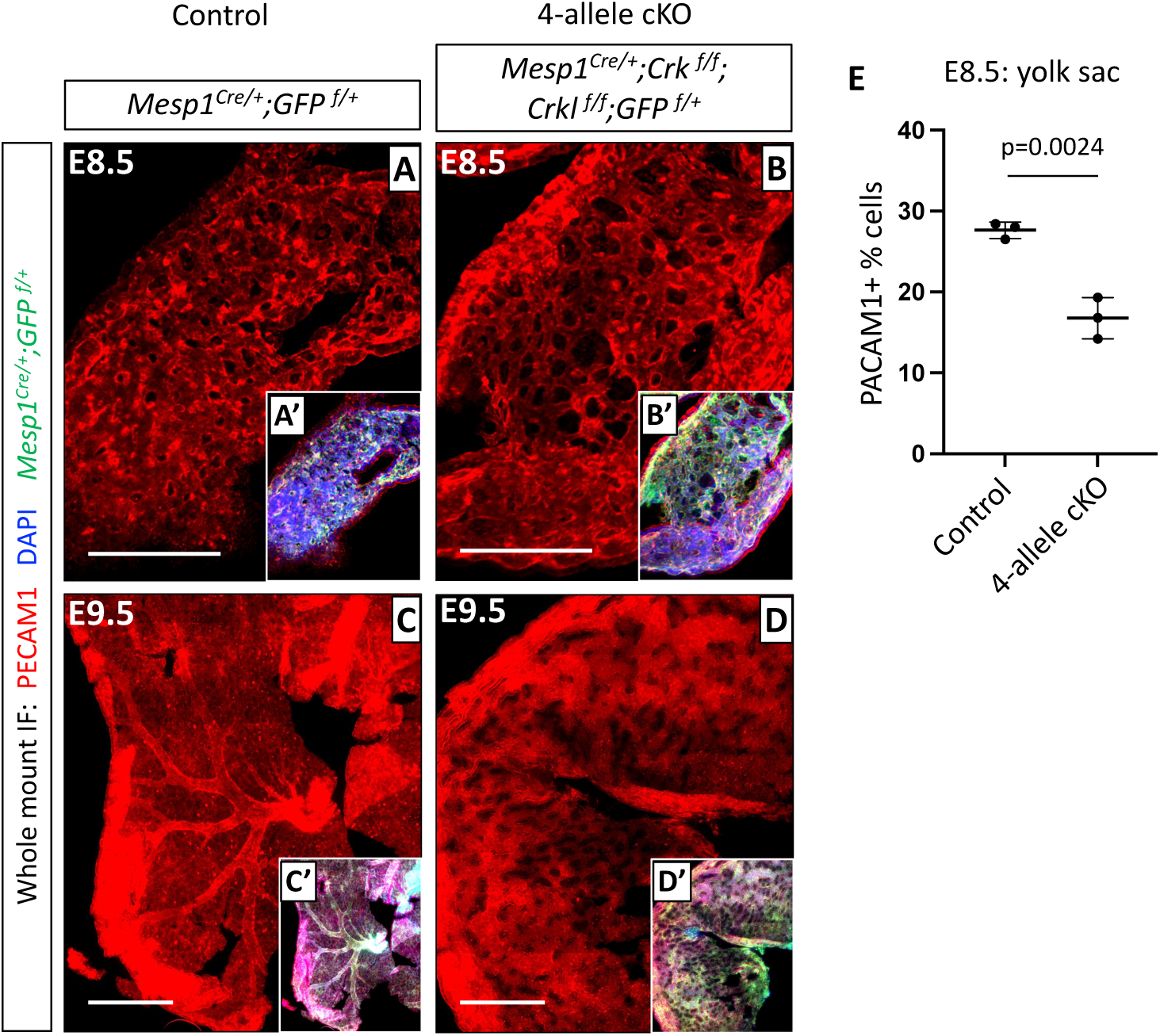
Loss of *Crk* and *Crkl* in the *Mesp1* lineage resulted in a disorganized vascular network within the yolk sac. (A-D) Whole mount immunofluorescence staining of anti-PECAM1 and anti-GFP antibodies in the yolk sac showing the vasculature network in control (*Mesp1^Cre/+^;GFP ^f/+^*) and 4-allele cKO (*Mesp1^Cre/+^;Crk ^f/f^;Crkl ^f/f^;GFP ^f/+^*) embryos at E8.5 and E9.5. There are less dense capillary plexuses in 4-allele cKO embryo at E8.5 (B), and the failure of vasculature remodeling (D) versus the normally remodeled vasculature in controls at E9.5 (C). Minimal N=3 embryos for each genotype and stage. Scale bar in (A, B): 250 μm. Scale bar in (C, D): 500 μm. (E) Quantification of the percentages of ECs (PECAM1+) in the yolk sac of the control (Cre negative) and 4-allele cKO embryos at E8.5 by flow cytometry. The graphs were plotted with mean and standard deviation by Prism 9 software. Each dot in the graph represents one embryo. Two-tailed Student’s t-test was used to evaluate statistical significance.

Next, we examined the vasculature in embryos themselves taking the same approach. We found that the blood vessels were severely disorganized in the 4-allele cKO embryos, versus controls at both E8.5 and E9.5 (Fig. 2; minimal N=3 for each genotype and stage). The vascular network in the head, heart, dorsal aorta and intersomitic vessels were all poorly remodeled in 4-allele cKO embryos (Fig. 2C-D) versus controls (Fig. 2A-B). Further, we also found that both genotypes of 3-allele cKO embryos exhibited similar vascular defects in the yolk sac and embryo as compared to the 4-allele cKO embryos, while 2-allele cKO embryos had normal vascular development (Fig. S3; minimal N=3 for each genotype). Therefore, our data suggest that *Crk* and *Crkl* are required in the *Mesp1* lineage for normal vascular formation in the yolk sac and embryos and that one allele of *Crk* or *Crkl* in *Mesp1* lineage is not sufficient to maintain normal vascular development in early mouse embryonic development.

**Figure 2.**
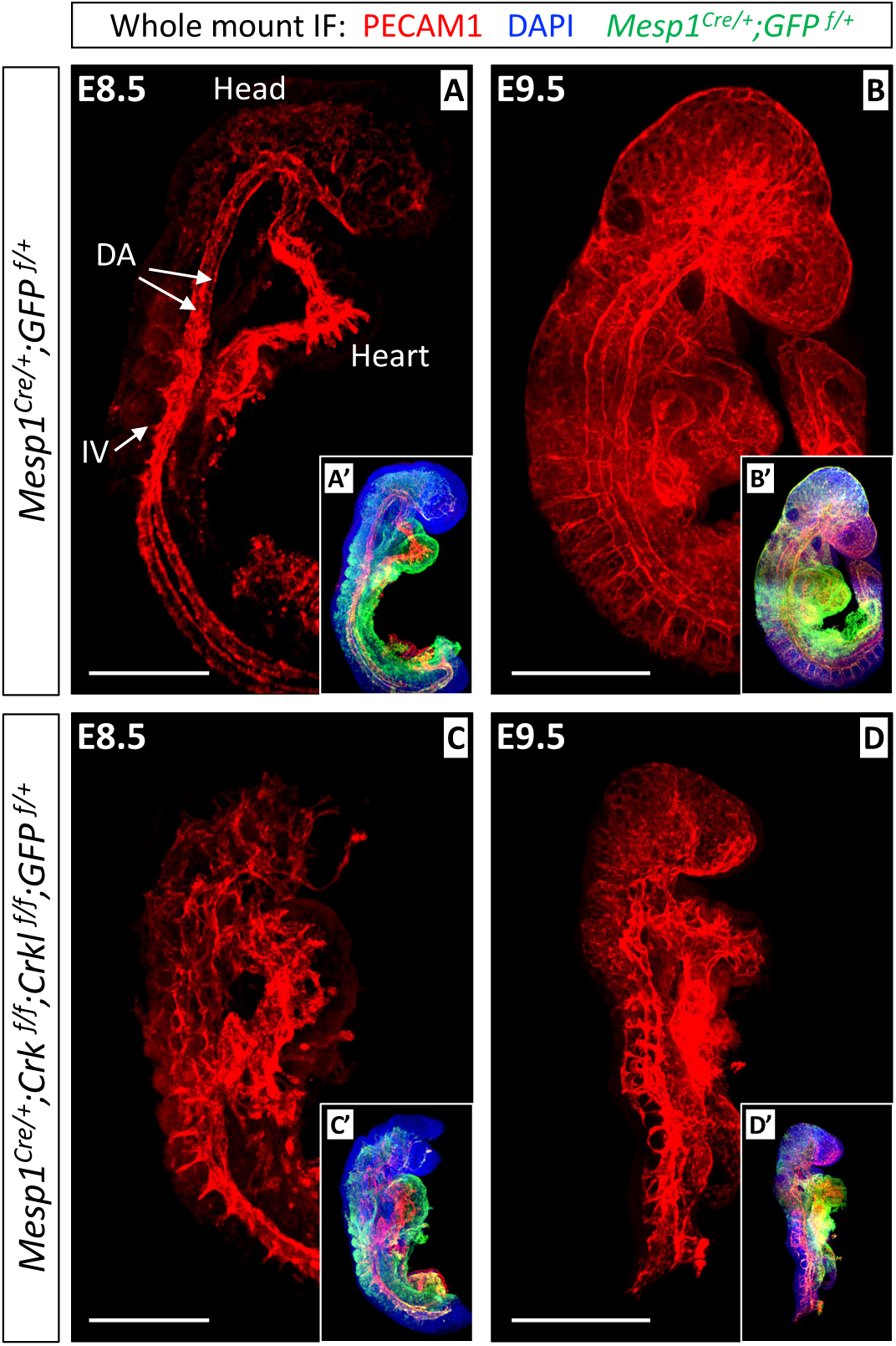
Loss of *Crk* and *Crkl* in the *Mesp1* lineage caused a disorganized vascular network in embryos. Whole mount immunofluorescence staining of anti-PECAM1 and anti-GFP antibodies in embryos shows the vasculature network in control (*Mesp1^Cre/+^;GFP ^f/+^*) and 4-allele cKO (*Mesp1^Cre/+^;Crk ^f/f^;Crkl ^f/f^;GFP ^f/+^*) embryos at E8.5 and E9.5. The blood vessels were severely disorganized in 4-allele cKO embryos at both E8.5 (C) and E9.5 (D). Insets depicts DAPI, and antibody staining (A’, B’, C’, D’). DA: dorsal aorta. IV: intersomitic vessels. Minimal N=3 for each genotype and stage. Scale bar in (A, C): 250 μm. Scale bar in (B, D): 500 μm.

Following this, we examined an earlier stage, E8.0, to investigate whether there were vasculogenesis defects in 4-allele cKO embryos. We performed whole mount RNAscope analysis of *Etv2* and *Pecam1* (Fig. 3A), and whole mount immunofluorescence of anti-VEGFR2 antibody (Fig. 3B) in control and 4-allele cKO embryos at E8.0 (minimal N=3 for each genotype). *Etv2* is expressed in immature vascular EC progenitors, while *Pecam1* is expressed in more mature and differentiated vascular ECs.^30,49^ VEGFR2 is expressed in all ECs.^31^ We found there were no gross changes in the expression pattern of *Etv2*, *Pecam1* and VEGFR2 in 4-allele cKO versus control embryos at E8.0, suggesting that vasculogenesis in the 4-allele cKO embryos is grossly normal (Fig. 3A-B). Thus, our data indicate that loss of *Crk* and *Crkl* in the *Mesp1* lineage does not affect vasculogenesis but suggest that defective angiogenesis might be responsible for the observed phenotype in later staged mutant embryos.

**Figure 3.**
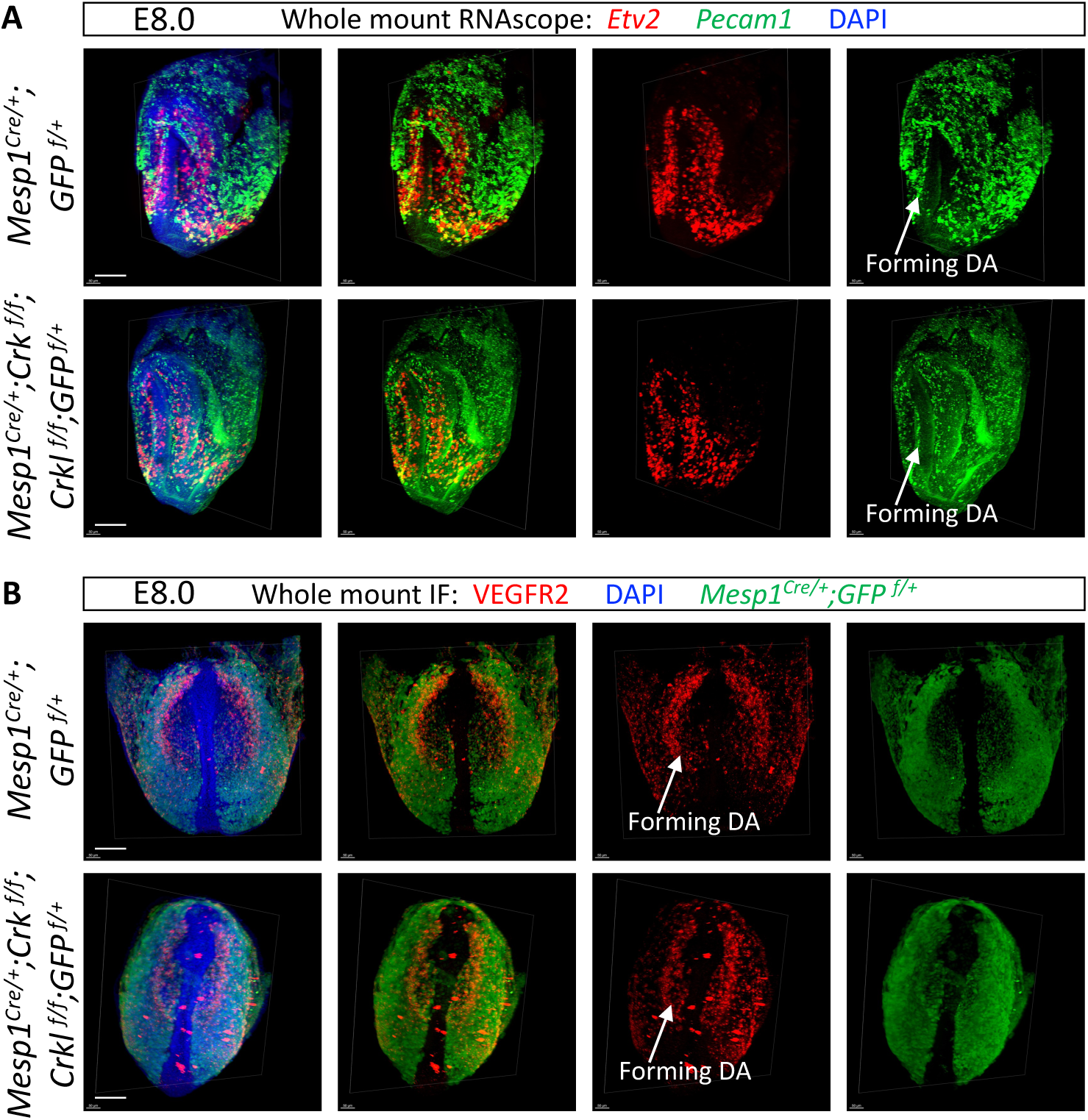
Loss of *Crk* and *Crkl* in the *Mesp1* lineage did not affect vasculogenesis at E8.0. (A) Whole mount RNAscope *in situ* hybridization staining of *Etv2* and *Pecam1* probes showing the EC progenitors and differentiated ECs, respectively, in the yolk sac and embryos in control (*Mesp1^Cre/+^;GFP ^f/+^*) and 4-allele cKO (*Mesp1^Cre/+^;Crk ^f/f^;Crkl ^f/f^;GFP ^f/+^*) embryos at E8. (B) Whole mount immunofluorescence staining of anti-VEGFR2 and anti-GFP antibodies showed the ECs in the yolk sac and embryos in the control and 4-allele cKO embryos at E8. Minimal N=3 for each genotype. Scale bar: 100 μm.

The 4-allele cKO embryos were slightly smaller than 2-allele controls at E8.5 but were dramatically smaller at E9.5 (Fig. 2C-D). Next, to investigate whether there were changes in cell proliferation and apoptosis in the 4-allele cKO embryos versus controls, which may partially contribute to the reduced size of embryos, we performed whole mount immunofluorescence staining of anti-phospho-histone H3 (pH3) antibody and TUNEL assays on embryos at E8.5 and E9.5 (minimal N=3 for each genotype and stage). We did not observe any gross changes of cell proliferation in the 4-allele cKO embryos versus controls at both E8.5 and E9.5 (Fig. S4). We also did not observe gross changes in apoptosis in the *Mesp1* lineage cells (Fig. S4). This suggests that the observed vascular defects might not result from changes in cell proliferation or apoptosis. Of interest, we observed a significant increase of apoptosis in non-*Mesp1* lineages in the dorsal side and frontal head regions of 4-allele cKO embryos versus controls at E9.5 but not E8.5 (Fig. S4), indicating that *Crk/Crkl* in the *Mesp1* lineage might function in a cell non-autonomous way to affect the survival of neighboring cells, or this could be a secondary phenotype due to hypoxia resulting from the disorganized vascular network with possible ineffective perfusion.

### Changes in cell clusters identified by scRNA-seq in Crk/Crkl mutants versus controls at E8.5

To better understand the underlying molecular mechanisms for defective angiogenesis, we performed scRNA-seq of the *Mesp1^Cre^* mesoderm lineage purified from 4- versus 2-allele cKO embryos at E8.5, when the vascular phenotype began to appear. The 2-allele cKO embryos were used as controls since they had normal vascular development (Fig. S3) and can be collected from the same litters with the 4-allele cKO embryos. For simplicity, the 2- and 4-allele cKO embryos used for the scRNA-seq are noted as control and cKO, respectively. The rostral half of embryos were micro-dissected and pooled, and GFP positive mesodermal lineage cells were purified for scRNA-seq by flow cytometry (Fig. 4A, Table S2). A total of 27,201 mesodermal lineage cells were sequenced from two replicates of control and cKO embryos, with the mean sequencing reads per cell ranging from 63,057 to 117,616 megabases (Table S2). The Seurat package ^43^ was used for quality control filtering of the single scRNA-seq dataset (Fig. S5), and Robust Integration of scRNA-seq (RISC) package ^44^ was used to integrate the control and cKO sequencing datasets and for identification of cell cluster types.

**Figure 4.**
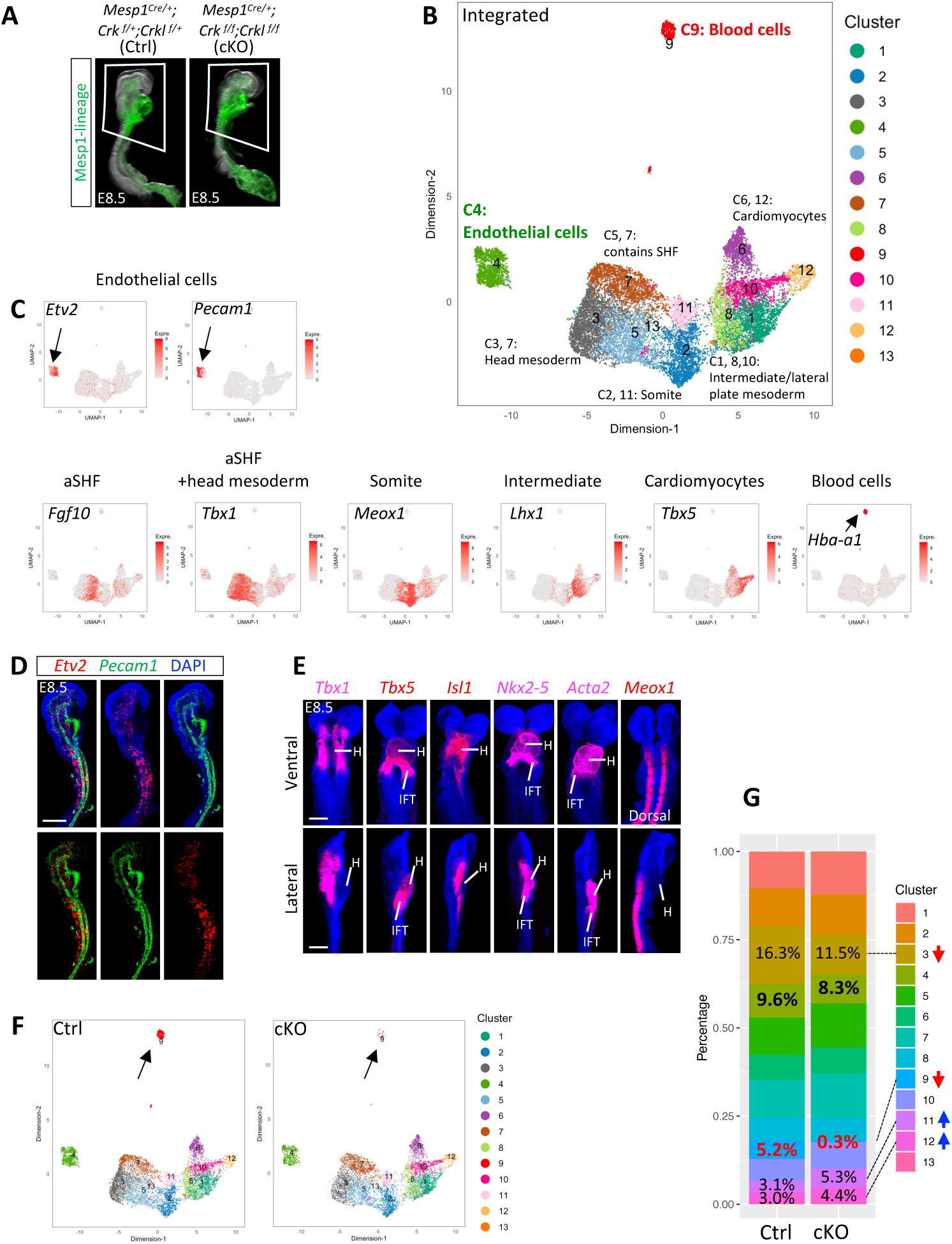
Single cell RNA-seq of mesodermal lineage cells at E8.5 in cKO versus control embryos. (A) Whole mount view of 2-(control, Ctrl) and 4-allele cKO (cKO) embryos at E8.5 shows the *Mesp1^Cre/+^;GFP ^f/+^* genetic lineage. The rostral part of embryos (indicated in the white quadrilateral) was micro-dissected and then GFP positive cells were sorted for scRNA-seq. (B) RISC Uniform Manifold Approximation and Projection (UMAP) shows the integrated control and cKO scRNA-seq datasets with cluster annotations. (C) UMAP plots using RISC software of the expression of selected maker genes used for identifying cell types indicated in (B). (D, E) Whole mount RNAscope *in situ* hybridization of *Mesp1^Cre/+^;GFP ^f/+^* embryos at E8.5 with probes stained for *Etv2*, *Pecam1*, *Tbx1*, *Tbx5*, *Isl1*, *Nkx2-5*, *Acta2*, and *Meox1* separately and/or together. Both Ventral and lateral views of embryos were shown in (E). Dorsal view of embryo was shown for *Meox1* in (E). N=3 for each probe in (D, E). aSHF: anterior second heart field. IFT: inflow tract. H: heart. Scale bar in (D, E): 200 μm. (F) UMAP plot using RISC shows the separated control and cKO scRNA-seq datasets with cluster identification. (G) Stack bar graph shows the proportions of mesodermal cells in each cluster divided by total mesodermal cells analyzed in either control or cKO sample. Note there was a slight decrease of proportion of mesodermal cells in cluster C3, significant decrease in cluster C9, and slight increase in cluster C11 and C12 in the cKO sample compared to the control. Two proportion Z test was performed to evaluate the statistically significant differences of proportions between control and cKO samples. No statistical significance of the change of the EC proportions between control and cKO samples. P values for cluster C3, C9, C11, C12 are < 2.20E-16, < 2.20E-16, 1.20E-14, 1.38E-07, respectively. Red and blue arrows in (G) indicate the decrease and increase of proportions, respectively.

We identified 13 clusters in the integrated datasets consisting of seven major cell types (Fig. 4B). The genes identified in each cell cluster are provided in the Supplementary Data 1. The identity of the cell type within each cluster is based upon the expression of marker genes (Fig. 4C, Fig. S6). The ECs are important for vascular development and we found that cluster C4 contains ECs as determined by expression of well-known EC specific marker genes including *Etv2*, *Pecam1*, *Cdh5* and *Tie1* (Fig. 4C, Fig. S6).^30,49,50^ We identified genes expressed in the anterior second heart field (aSHF; Fig. 4C, Fig. S6) that forms the cardiac outflow tract, head mesoderm/mesenchyme that contributes to connective tissue, mesoderm of the somites, intermediate and lateral plate mesoderm and cardiomyocyte progenitor cells (Fig. 4C, Fig. S6). Cluster C9 contains blood cells which express blood cell specific genes such as *Hba-a1*, *Gypa* and *Snca* (Fig. 4C, Fig. S6). Spatial expression of selected marker genes, such as *Etv2* and *Pecam1* for ECs, *Tbx1*, *Tbx5*, *Isl1*, *Nkx2-5* and *Acta2* for the SHF and/or heart, and *Meox1* for somites, in *Mesp1^Cre/+^;GFP ^f/+^* control embryos at E8.5 were confirmed by whole mount RNAscope *in situ* hybridization (Fig. 4D-E).

We found that there were no cell clusters that were completely missing in the cKO versus control scRNA-seq datasets (Fig. 4F, Fig. S7). There was no significant change of the proportion of ECs compared to the total GFP positive cells (Fig. 4G, Fig. S7), suggesting normal numbers of EC cell types. Interestingly, there was a significantly decreased proportion of blood cells in the mutant embryos (Fig. 4F-G, Fig. S7). As mentioned above, PECAM1 positive cells in the yolk sac, which contains hemogenic ECs that produce hematopoietic progenitor cells (HPCs) entering the circulation after the heart starts beating^51–56^, were significantly reduced in the yolk sac of 4-allele cKO versus control embryos at E8.5 (Fig. 1E). We quantified percentages of HPCs (PECAM1+cKit+^52^) and found the significant decrease of HPCs in the yolk sac of 4- and 3-allele cKO embryos compared to the controls (Fig. S8). Our data suggest that the observed reduced blood cell proportion in the scRNA-seq dataset may be secondary due to defective vascular development. In addition, decreased cell proportions were observed in the head mesoderm/mesenchyme, and there were slightly increased numbers of cardiomyocytes and somitic cells as compared to the total number of GFP positive mesodermal cells in cKO versus control scRNA-seq datasets (Fig. 4G, Fig. S7). Therefore, the scRNA-seq data suggest that inactivation of *Crk/Crkl* might affect cell proportions, but it doesn’t result in complete absence of any population of *Mesp1* lineage cells. However, to understand changes in cell states, we examined gene expression level changes in ECs.

### Upregulation of genes crucial for angiogenesis and vasculature development in ECs in cKO versus control embryos at E8.5

Next, we analyzed scRNA-seq data of ECs because these cells are related to the observed vascular phenotype seen in cKO embryos. Differentially expressed genes (DEGs; adjust p value < 0.05, |log_e_^(fold change)^| > 0.25) in ECs from the cKO versus control scRNA-seq datasets were calculated using RISC software,^44^ and the full list of DEGs is shown in the Supplementary Data 2. There were 334 up-regulated and 113 down-regulated DEGs identified in the EC population (cluster C4) in cKO versus control embryos (Fig. 5A-B). Significantly more up- versus down-regulated DEGs identified in ECs (p<2.2E-16) suggests that *Crk/Crkl* may normally act in a signaling pathway to modulate EC gene expression. We also noticed that the overall relative fold changes of DEGs were relatively small with most of them were in the range of |log_e_^(fold change)^| < 1, but the adjusted p values are highly significant (Fig. 5A). This is not surprising since CRK/CRKL are adaptor proteins that mediate multiple signal transduction pathways and therefore indirectly regulate expression of downstream genes in a collective way, in addition to their prominent roles in mediating posttranslational modification of protein activities.^57^

**Figure 5.**
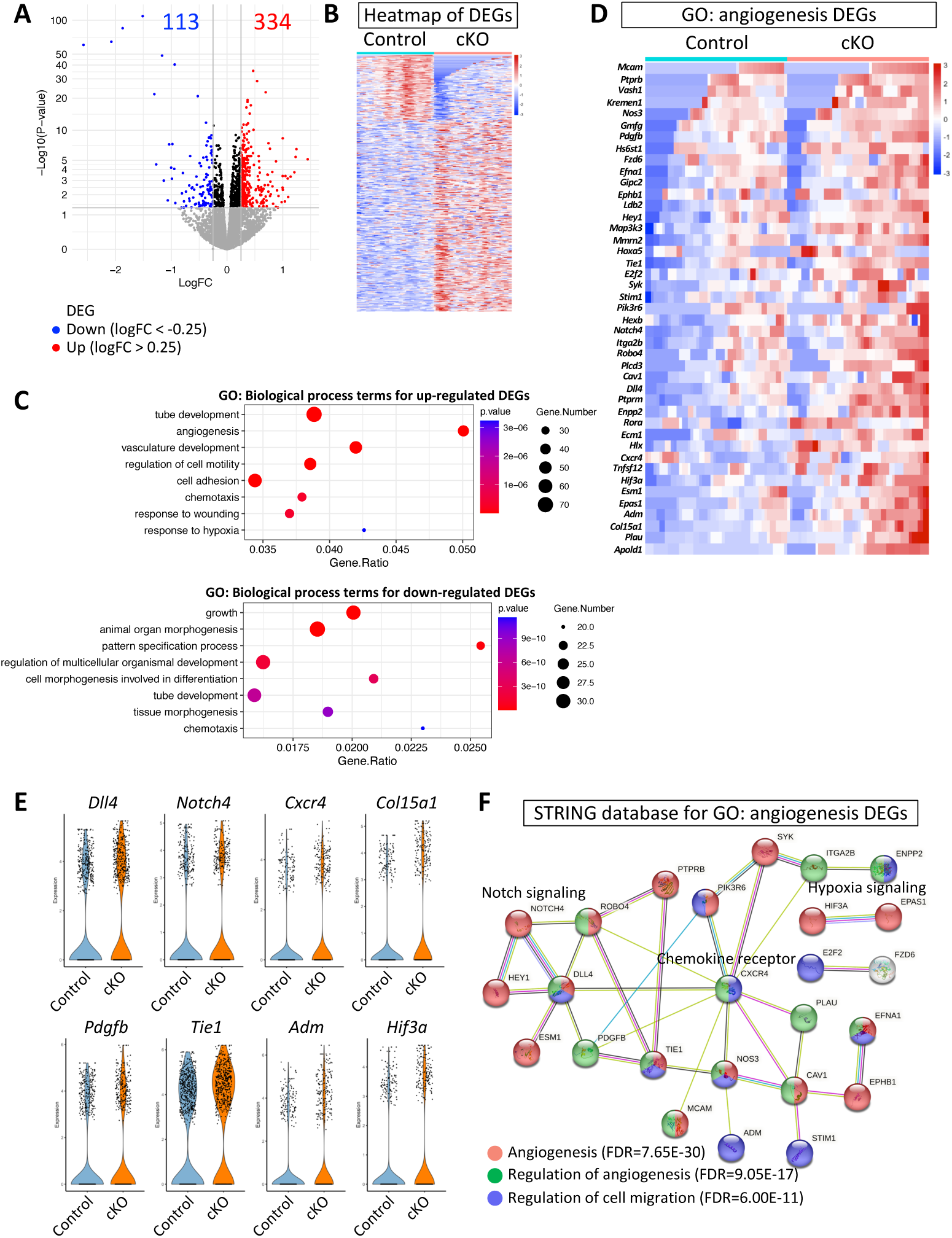
A significant number of angiogenesis and cell adhesion/migration genes were up-regulated in the EC population in the cKO versus control scRNA-seq datasets. (A) A volcano plot shows the differentially expressed genes (DEGs) with log_e_(Fold change) (x axis) and - log_10_(adjusted p value) (y axis). Red and blue dots represent DEGs with log_e_(Fold change) > 0.25 & adjusted p value < 0.05, log_e_(Fold change) < -0.25 & adjusted p value < 0.05, respectively. Numbers of up- and down-regulated DEGs in cKO versus control ECs are indicated in red and blue font, respectively. (B) A heatmap shows the expression of all detected DEGs in the control and cKO ECs. (C) Gene Ontology (GO) analysis of Biological Process terms for up- and down-regulated DEGs identified in ECs. Top Biological Process GO terms with smallest, most significant, p value were selected for the plots. Gene.Number in the side bar shows the number of DEGs detected per GO term. Gene.Ratio in the x-axis is the ratio of number of DEGs in the GO term with respect to the number of all the genes in that GO term. (D) A heatmap shows the expression of DEGs selected from the angiogenesis Biological Process GO term in (C) in the control and cKO samples in ECs. (E) Violin plots generated using RISC software shows the expression of selected DEGs from angiogenesis Biological Process GO term in (C) in the control and cKO samples in ECs. (F) STRING protein interactions analysis for DEGs selected from angiogenesis Biological Process GO term in (C).

To investigate the types of DEGs found in ECs, we performed gene ontology (GO) analysis using ToppFun software within the ToppGene Suite.^45^ The full list of GO analysis of DEGs for Molecular Function, Biological Processes and Pathway GO terms are shown in the Supplementary Data 3. Biological Process GO gene sets had the most significant p-value (Supplementary Data 3). Dot-plots of selected top GO terms are shown in Fig. 5C. The up-regulated DEGs were enriched in two major categories of Biological Process GO terms including angiogenesis/vasculature development and cell adhesion/migration, while the down-regulated DEGs were enriched in multiple organ development/morphogenesis GO terms (Fig. 5C, Fig. S9). Among the up-regulated DEGs, many overlapped such as there were 69 genes shared among the two main categories of angiogenesis/vasculature and cell adhesion/migration among 155 genes (Fig. S10; calculated in BioVenn website^58^). This reflects the fact that angiogenesis requires cell migration (migration/adhesion) functions. We also performed a GO analysis of Biological Processes in the STRING database and the results are similar (Supplementary Data 4). Multiple angiogenesis and cell migration genes were up-regulated in the ECs in the cKO versus control embryos, such as NOTCH signaling genes *Dll4* ^32–34,36,38,59^, *Hey1* ^37^ and *Notch4* ^60,61^, chemokine receptor gene *Cxcr4* ^62–64^, collagen gene *Col15a1* ^65,66^, hypoxia signaling genes *Epas1* ^67,68^, *Hif3a* ^69^ and *Ddit4* ^70^, growth factor gene *Pdgfb* ^71^, and preprohormone gene *Adm* ^72–75^ (Fig. 5D-F). Overexpression of *Dll4* ^32^ and *Cxcr4* ^62^ in mouse ECs has been shown to cause defective vessel branching or angiogenesis. We found upregulation of multiple genes important for regulating cell migration, such as focal adhesion genes *Itga2*, *Itga2b*, *Lama4* and *Mcam* (Fig. S9-10). Many of those up-regulated DEGs promote angiogenesis, whereas some promote or restrict cell migration (Fig. S10), suggesting that a combination of these changes can lead to dysregulated angiogenesis. Those that were down-regulated were general developmental genes but they did not provide particular insights as to connections to the observed vascular phenotype, although they could be secondary to the vascular defect or a primary defect. In conclusion, the initial analysis of the total EC cluster suggests that inactivation of *Crk/Crkl* leads to angiogenesis defects. To further define the mechanism, we delved more deeply into the scRNA-seq data.

### EC population subcluster analysis showed significantly increased differentiated angiogenic cells in cKO embryos versus controls at E8.5

To define the differences of ECs more specifically between cKO versus control embryos, we performed subcluster analysis of the EC population. At E8.5, ECs are not yet specified to subtypes, such as arterial or venous ECs, but they are nevertheless heterogenous in cellular states (Fig. 6A). *Etv2* is a master regulator gene driving EC specification, while *Cdh5* is a direct transcriptional target of ETV2.^76,77^ *Pecam1* marks most early ECs and also functions downstream of *Etv2*.^50^ Therefore expression of genes including *Cdh5* and *Pecam1* mark early steps of maturation of ECs downstream of *Etv2*.^30,77^ We identified subclusters including immature EC progenitors versus more mature ECs based on the gradient expression of *Etv2*, *Cdh5* and *Pecam1* (Fig. 6A-C).^30,77^ We identified angiogenic ECs, comprising a subset of those expressing *Cdh5* and *Pecam1*, based upon expression of *Apln*, *Dll4* and *Hey1* (Fig. 6A-C).^78,79^ The full lists of the marker genes used to identify subclusters are shown in the Supplementary Data 5. There were no missing subclusters and cell subtypes; however, there was significantly increased cell proportion (2 fold increase in proportion) in subcluster C2, which contains cells at a more differentiated state, which are undergoing angiogenesis, and decreased cell proportion in subclusters C3, C4, C8 which contain *Etv2* expressing EC progenitors in cKO versus control embryos (Fig. 5D-E). Based upon expression of marker genes such as *Dll4*, *Cxcr4*, *Pdgfb, Unc5b* and *Cxcr4*,^31^ this subcluster may contain angiogenic tip cells and these genes were significantly increased in expression in cKO versus control ECs (Fig. 6G; Fig. S11). Cell cycle calculations for each subcluster showed that cells in subcluster C2 are overall less proliferative compared to other subclusters (Fig. S12), which is consistent with a characteristic of tip cells of having reduced proliferative ability.^31^ Further, there was a decreased proportion (reduced by half) of cells in the G2M cell cycle phase and increased proportion of cells in S and G1 cell cycle phases in subcluster C2 in cKO embryos, supporting the idea that there are more mature angiogenic cells, possibly tip cells, which are less proliferative, that are either undergoing DNA replication or arrest at G1 (Fig. 6F). Expression of selected genes in the Fig. 6B in ECs of control and cKO embryos were shown in Fig. 6G. Among the six genes, *Dll4* and *Hey1* were identified as up-regulated DEGs and they are expressed in more cells as shown in Fig. 6G and Supplementary Data 2. Some other genes were not significantly changed in expression but were expressed similarly in more (*Apln, Cdh5, Pecam1*) and less (*Etv2*) cells, respectively (Fig. 6G). As *Dll4* and *Hey1* are two key NOTCH pathway genes involved in angiogenesis, arterial fate specification, and vessel remodeling,^37,38^ these observations support the suggestion that *Crk/Crkl* regulate early embryonic vessel formation in part through NOTCH signaling.

**Figure 6.**
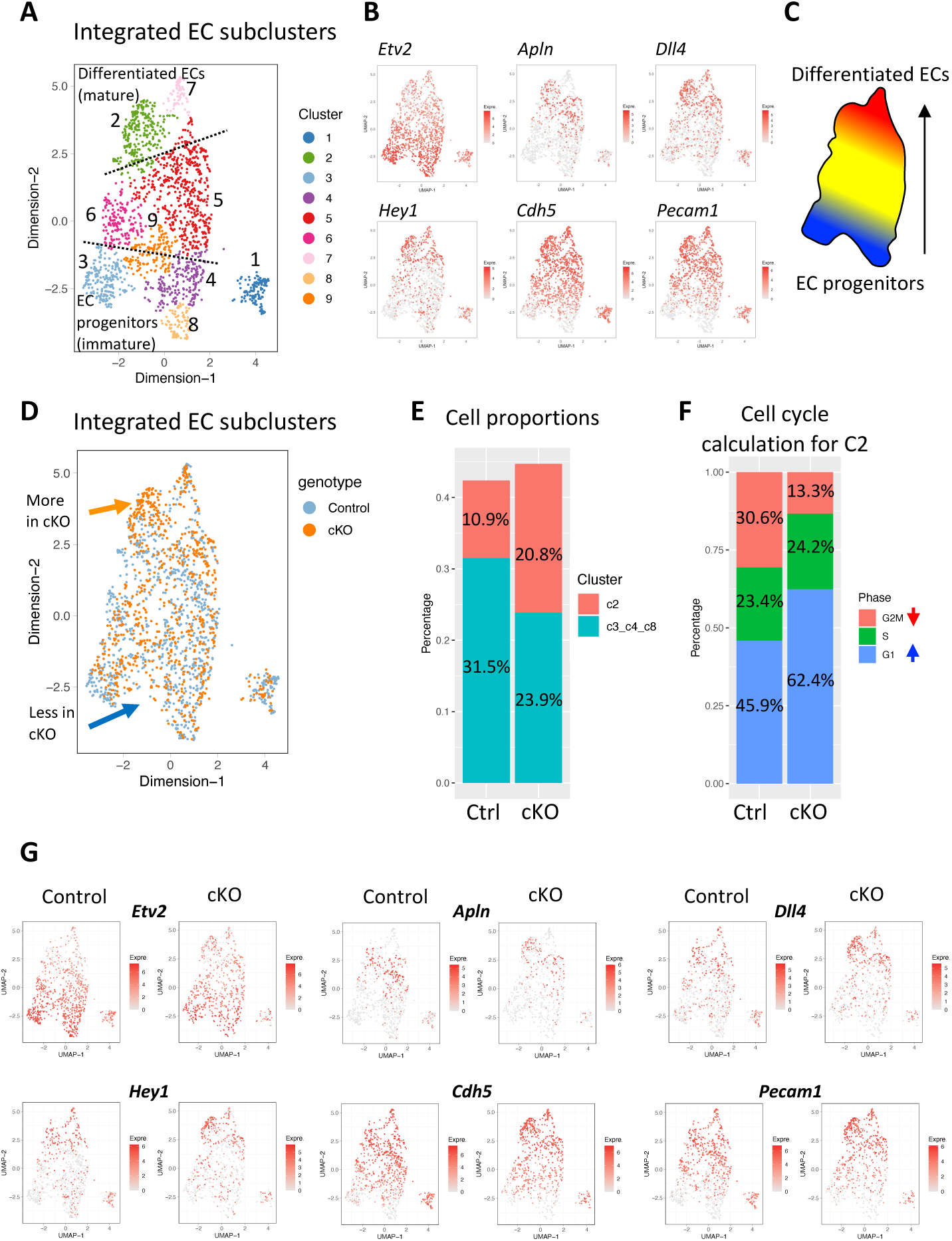
Loss of *Crk/Crkl* resulted in a significant increase of angiogenic ECs. (A) UMAP using RISC software shows the integrated EC subclusters obtained from the EC population, C4 in Fig. 4B. (B) UMAP plots show the expression of selected maker genes used for identifying the EC subpopulations. *Etv2* marks the immature, EC progenitors. *Apln*, *Dll4* and *Hey1* mark the angiogenic ECs. *Cdh5* and *Pecam1* mark the mature and more differentiated ECs. (C) A cartoon shows the gradient distribution of EC subpopulations from progenitors to more differentiated ECs in (A). (D) UMAP shows the ECs in integrated control (blue) and cKO ECs (orange). Orange and blue arrows in (D) indicate the increased and decreased ECs in cKO versus control embryos, respectively. (E) Stack bar graph shows the proportions of EC numbers in selected subclusters divided by total EC numbers analyzed in either control or cKO samples. (F) Stack bar graph shows the proportions of EC numbers in the cell cycle of G2M, S and G1 phases divided by the total EC numbers in subcluster C2 in either control or cKO sample. Two proportion Z test was performed to identify statistically significant differences of proportions between control and cKO samples in (E, F). P value for the C2 and C3_C4_C8 in (E) is 6.67E-09 and 4.05E-04, respectively. P value for the G2M phase and G1 phase in (F) is 8.05E-04 and 9.9E-03, respectively. Red and blue arrows in (F) indicate the decrease and increase of proportion, respectively. (G) UMAP shows the expression of selected genes in (B) in ECs in cKO versus control scRNA-seq datasets.

### Expansion of Dll4 expressing angiogenic cells in cKO versus control embryos at E8.5

*Dll4* is highly expressed in the angiogenic tip cells and expressed at reduced level in the stalk cells, which play crucial role during sprouting angiogenesis, and expression of *Dll4* is crucial for arterial fate specification.^31,80^ Our *in vivo* whole mount RNAscope analysis of *Dll4* confirmed that it is expressed in more ECs in cKO embryos. We found that there was an expansion of *Dll4* expressing cells in the head, dorsal part of the embryo (with respect to the dorsal aorta), and somites of cKO embryos versus controls (Fig. 7A-B; N=3 for each genotype), supporting the bioinformatic results. Combined with the previous cell cycle analysis of cells in subcluster C2, which exhibited overall decreased cell proliferation, our data suggest that the observed disorganized pattern of vascular network in the cKO embryos is due to failed coalescence or remodeling which is supported by less mature arteries but more *Dll4+* angiogenic cells (Fig. 7A-B). Immunofluorescence staining on sections for anti-NOTCH1 intracellular domain (NICD) antibody was performed to evaluate further the expression changes of *Dll4* and the result showed similar or a slight expansion of NICD signal in the dorsal and somite regions of the cKO embryos (Fig. 7C-D) (N=3 for each genotype), which collaborated well with the *Dll4* and *Hey1* findings. Together, these observations further support a *Crk/Crkl*-dependent NOTCH signaling, in part, underlying early embryonic vessel formation through arterial remodeling.

**Figure 7.**
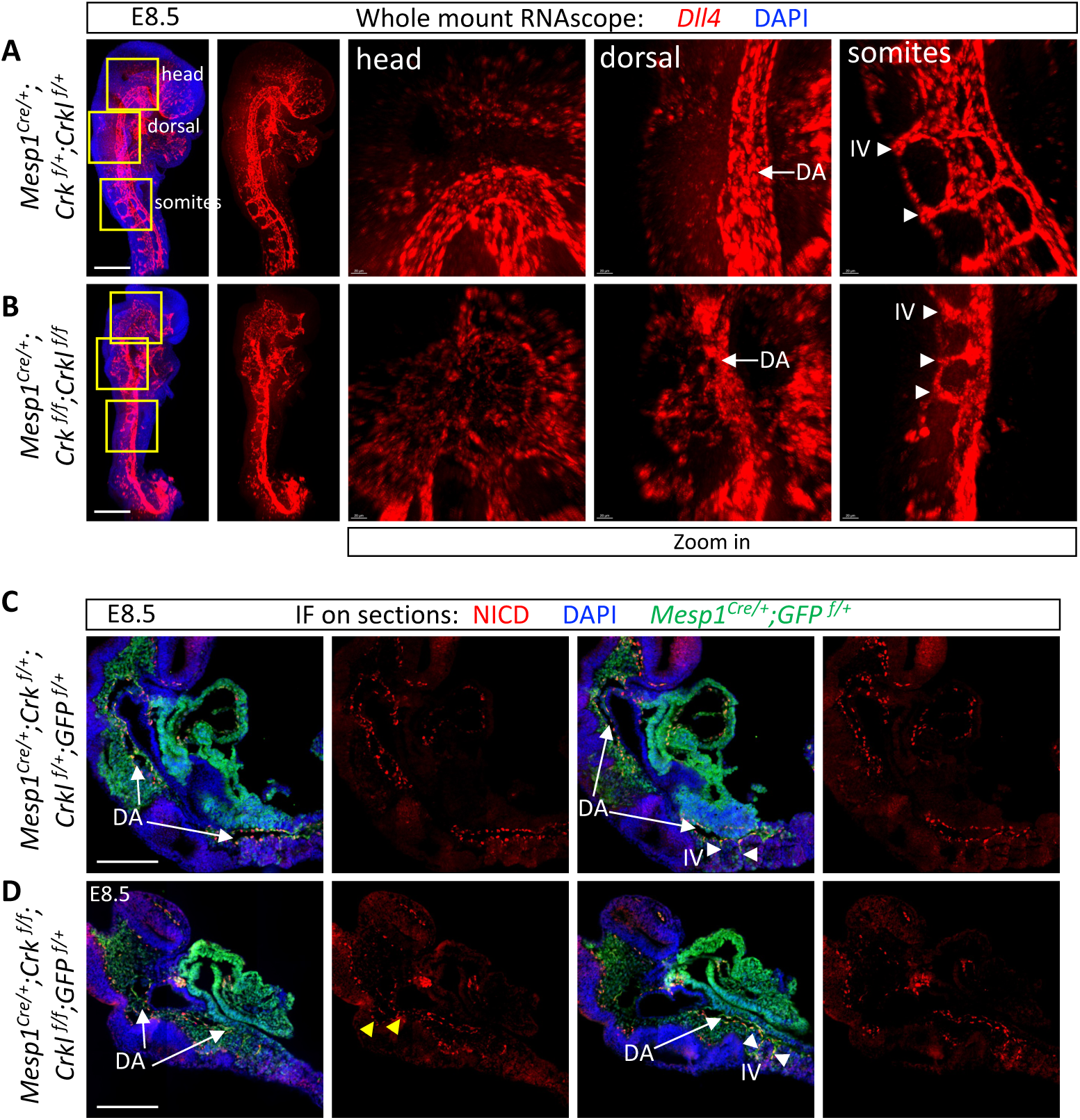
Increased angiogenic ECs in cKO embryos were identified using *Dll4* and NICD staining at E8.5. (A, B) Whole mount RNAscope of a *Dll4* probe in control (A) and cKO (B) embryos, with and without DAPI signal. Yellow boxes of the head, dorsal part and intersomitic vessels (A, B) were expanded in images on the right. (C, D) Immunofluorescence staining of anti-Notch1 intracellular domain (NICD) and anti-GFP antibodies on paraffin sections of control (C) and cKO (D) embryos. Thickness of sections in (C, D): 10 μm. Minimal N=3 for each genotype. DA: dorsal aorta (white arrow with tail). IV: intersomitic vessel (white arrow without tail). Scale bar in (A-D): 200 μm.

### Accelerated EC differentiation in cKO versus control embryos at E8.5

To explain why we observed relatively fewer progenitor and more of mature, and perhaps atypical angiogenic ECs in cKO embryos versus controls, we bioinformatically analyzed the EC cell fate progression and dynamics of the scRNA-seq data at E8.5 by using scVelo^47^ and CellRank^48^ software. Initial (progenitor) and terminal (differentiated) states of ECs were identified along with pseudo developmental time, also known as latent time. This takes advantage of cell state heterogeneity within a population with data from a single developmental time point. We found that ECs could differentiate normally from immature, progenitor cells to more mature, differentiated cells in cKO and control embryos (Fig. 8A; Fig. S13), suggesting that there was not a failure in cell fate progression by inactivating *Crk/Crkl*. Next, we identified genes, referred as lineage genes, that were progressively increased in expression during EC differentiation from immature to mature ECs for each scRNA-seq sample. We generated a cell fate probabilities heatmap of those genes at different developmental states, using one of the control samples (Fig. 8B; Supplementary Data 6). To better understand the changes of transcriptional gene expression signatures at different maturation status during EC differentiation, we manually divided the genes into four groups based on the time of their serial activation and performed GO analysis (Fig. 8C-D). Our analysis showed that the earlier and late activated genes were enriched into distinct GO terms (Fig. 8C-D). The late activated genes were highly enriched in angiogenesis and cell migration related Biological Process GO terms and extracellular matrix, focal adhesion Cellular Component GO terms (Fig. 8C-D).

**Figure 8.**
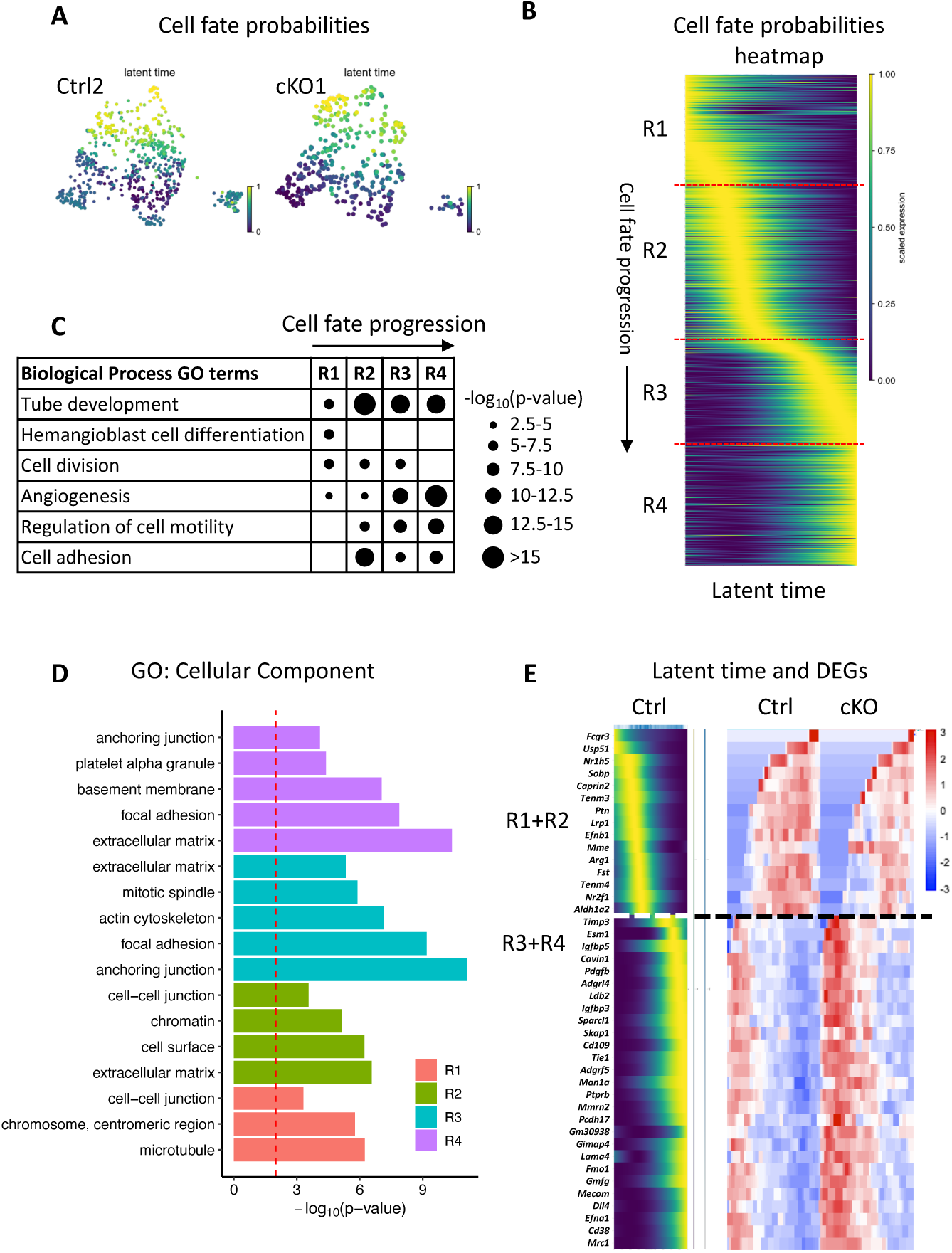
Dysregulation of genes promoting EC angiogenesis/differentiation occurred in cKO embryos at E8.5. (A) UMAP plot using CellRank software shows the cell fate probabilities of ECs differentiating to mature ECs in Ctrl2 and cKO1 samples. (B) Heatmap using CellRank software shows the expression of lineage genes whose expression correlates with the transition of ECs from progenitor to more differentiated cells which is indicated as latent time in the Ctrl2 sample. The full list of genes is shown in Supplementary data 6. The lineage genes were manually divided into 4 groups, including R1, R2, R3 and R4, based on their expression window during latent time. (C) Selected Biological Process GO terms analysis of 4 groups of lineage genes in (B). Dot sizes indicate distinct p values. (D) Selected Cellular Component GO analysis of 4 groups of lineage genes in (B). (E) Heatmap using CellRank (left) and RISC (right) shows the expression of the overlapping lineage genes among 4 scRNA-seq datasets and DEGs between control and cKO samples, respectively. White/black dashed line divides the early and late activated lineage genes.

We then analyzed changes of lineage genes in ECs between cKO and control embryos to investigate whether there is a change in any aspect of cell fate progression during EC differentiation. Among a total of 255 overlapping genes in both replicates between cKO and control datasets, we identified that 42 of them are DEGs including 28 up-regulated and 14 down-regulated genes (p value < 2.2E-16; Fig. 8E). Almost all the earlier expressed lineage genes among the 42, were down-regulated and all late expressed lineage genes among the 42, were up-regulated in ECs in cKO versus control embryos (Fig. 8E). Therefore, our data suggests that increased expression of genes for an atypical, differentiated state at later time points may result in precocious angiogenesis that fails to coalesce or remodel to produce functional vessels.

## Discussion

In this report, we found that inactivation of *Crk* and *Crkl* in the *Mesp1* lineage disrupted early mouse blood vessel development in both the yolk sac and embryo proper. Mutant embryos died in early gestation with severely disorganized blood vessels. Single cell transcriptomic analysis and *in vivo* validation revealed the presence of enhanced angiogenesis of ECs when both *Crk/Crkl* were inactivated together in the *Mesp1* lineage.

New blood vessels are formed through vasculogenesis and then angiogenesis during embryogenesis.^29,81^ In the mouse, vasculogenesis occurs before heartbeat begins (E8.0-E8.25) and angiogenesis to create more branches and remodel those branches, occurs next in the yolk sac and embryo proper.^27–30^ *Mesp1* begins to be expressed when the mesoderm forms at gastrulation (E6.5),^41^ thus inactivation of *Crk/Crkl* using *Mesp1^Cre^* begins to occur during gastrulation and might affect both vasculogenesis and angiogenesis. We show in the model in Fig. 9A, that *Crk/Crkl* have a specific function in angiogenesis rather than vasculogenesis. Therefore, we suggest that *Crk/Crkl* in the mesoderm are dispensable for vasculogenesis. We observed enhanced and disorganized blood vessels, specifically in the early arteries, which were identified by *Pecam1* and *Dll4* expression in embryos (Fig. 9A).

**Figure 9.**
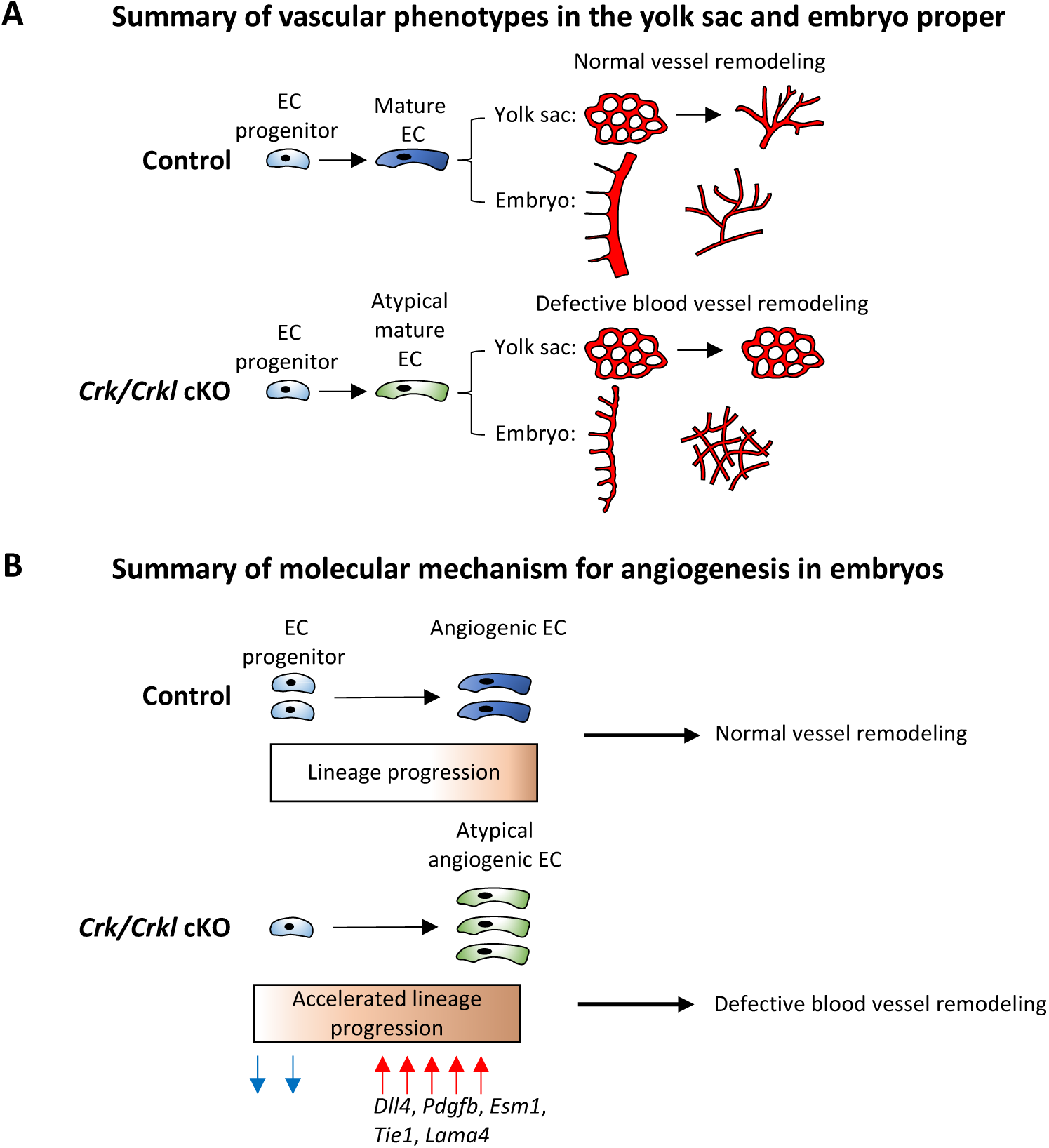
Schematic models of the mechanisms by which *Crk* and *Crkl* function in blood vessel development in the early embryonic mesoderm. (A) Schematic model shows the vascular phenotypes in the yolk sac in control and *Crk/Crkl* cKO (*Mesp1* lineage) embryos during early embryonic vascular development. Defective blood vessel remodeling is illustrated in cKO embryos. Abnormal ECs are indicated as green color in cKO embryos versus blue color in controls. (B) Schematic model shows a molecular mechanism leading to abnormal angiogenesis in cKO embryos. In brief, loss of mesodermal *Crk/Crkl* results in an increased number of atypical differentiated angiogenic ECs (green color versus blue in controls) due to accelerated lineage progression (orange color in box) during EC maturation. Decrease of expression of genes expressed in EC progenitor cells (blue, down arrows) and increase of expression in angiogenic, more mature ECs (red, up arrows) are shown. Selected genes are indicated below the red arrows.

We performed scRNA-seq of mesodermal cells to explore the molecular mechanism for the observed vascular defects upon inactivation of both *Crk/Crkl* (cKO). We focused on EC population since those cells are more related to the observed vascular phenotype in mutant embryos. We found that ECs are surprisingly heterogenous in composition even at this early developmental stage (E8.5). This made it possible to assess changes in EC composition and maturation. Using bioinformatic approaches, we found there was a relative decrease in *Etv2* expressing EC progenitor cells and increase of more differentiated angiogenic ECs, which was further confirmed *in vivo* showing increased amount of *Dll4* expressed cells, in cKO versus control embryos (Fig. 9B). This may be due to precocious and atypical cell fate progression during EC differentiation (Fig. 9B). Further bioinformatic analysis suggested that the dysregulation of these genes that affect EC function may contribute to accelerated cell lineage progression (Fig. 9B). Combined with cell cycle analysis for those more angiogenic ECs, we suggest this causes failed vascular coalescence and remodeling in the cKO embryos (Fig. 9B). Next, we want to discuss which signaling pathways might be mediated by CRK/CRKL during early angiogenesis process.

It has been well studied that VEGF signaling plays crucial roles in both vasculogenesis and angiogenesis.^40^ *In vitro* studies using primary human umbilical vein ECs showed that CRK acts downstream of VEGF signaling for regulating multiple EC behaviors including cell migration, survival and proliferation via posttranslational modification.^82,83^ CRK mediates VEGFR2 signaling for cell migration via activating integrin, regulating focal adhesion formation and modulating actin dynamics.^83^ CRK also mediates VEGFR3 signaling needed for EC proliferation, migration and survival through activating various downstream signaling pathways including ERK, AKT and JNK.^82^ Thus, VEGF signaling via loss of *Crk/Crkl* is likely affected, resulting in altered downstream effects. Other adaptor proteins, such as GRB2, FRS2 and Nck, also have been reported to mediate VEGF signaling during EC development.^82,83^ It is possible that some of the vascular defects we observed are due to improper signaling mediated by other adaptors that are present in the cells.

NOTCH signaling has been widely studied in mouse, zebrafish and human for its crucial roles in regulating angiogenesis, arterial/venous fate specification and vessel remodeling.^31,84^ In the mouse, genetic ablation of NOTCH signaling pathway genes including *Notch1* ^35^, *Notch4* ^85^, *Jag1* ^86^, *Hey1*, *Hey2* ^37^, as well as inactivation of one allele of *Dll4* ^36,38,87^ or both alleles of *Dll4* ^38^, or overexpression of *Dll4* ^32^ and *Notch4*^60^ all resulted in defective angiogenesis. Our scRNA-seq data showed increase in expression of several NOTCH signaling genes such as *Dll4*, *Notch4*, *Hey1* and *Hey2* in ECs in the cKO versus control embryos, which may explain or partially explain the observed angiogenesis defects in mutant embryos. Both scRNA-seq and our *in vivo* data indicated increased number *Dll4* positive angiogenic cells with overall reduced cell proliferation, suggesting that there is failed remodeling of blood vessels into mature arteries. CRK/CRKL have been well studied *in vitro* for their functions in regulating cellular cytoskeleton and multiple cellular behaviors including cell migration, proliferation, survival, and differentiation.^57,88,89^ CRK/CRKL might act to indirectly regulate NOTCH signaling via adjusting actin-based cytoskeleton assembly and affecting cell migration.^90–93^

VEGF and NOTCH signaling are tightly coupled to balance tip and stalk cell selection in regulating sprouting angiogenesis.^31^ *In vivo* and *in vitro* studies have indicated that VEGF-A signaling acts upstream of NOTCH signaling during angiogenesis by controlling *Dll4* expression, but how VEGF-A signaling regulates *Dll4* is less well understood.^31^ In turn, NOTCH signaling has been shown by *in vivo* and *in vitro* studies to control VEGF signaling during angiogenesis by regulating the expression of VEGF receptors.^31^ Thus, it is likely that loss of *Crk/Crkl* disrupts the balance of VEGF and NOTCH signaling in regulating angiogenesis, which results in the observed vascular defects.

In addition, our bioinformatic analysis of scRNA-seq data suggest that there is accelerated EC differentiation and defective vascular remodeling when inactivating *Crk/Crkl* in ECs. Endothelial tip cells are less proliferative and have highly motility; while the neighboring stalk cells need to proliferate and establish cell-cell adhesion, junctions and communication in order to promote vessel coalescence and remodeling to functional vascular network.^31^ Loss of *Crk/Crkl* in ECs caused up-regulation of multiple cell adhesion/migration genes in our data, which may affect tip/stalk cells functions that further contribute to the accelerated EC maturation and defective remodeling of vessels. Our data showed that the chemokine receptor gene, *Cxcr4* was significantly up-regulated in all ECs also in angiogenic ECs (subcluster C2). Studies in the mouse showed that loss of function of *Cxcr4* caused defective vascular development, while gain of function of *Cxcr4* in ECs resulted in defective vascular remodeling, which suggest a dosage sensitive requirement of *Cxcr4* in ECs for normal vascular development.^62–64^ Other cell adhesion/migration genes such as *Adm*, *Col15a1*, *Lama4*, *Mcam*, *Itga2* and *Itga2b*, were also up-regulated in ECs in the mutants. Malfunctions of preprohormone gene *Adm* ^72–75^ and collagen gene *Col15a1* ^65,66^ also caused angiogenesis and/or vascular defects. The dysregulation of cell adhesion/migration related genes in ECs may also contribute to the observed vascular defects.

In conclusion, we found that *Crk/Crkl* are required in the early mesoderm for normal angiogenesis both in the yolk sac and embryo. We used scRNA-seq to explore the altered transcriptome of mesodermal cells when the expression of *Crk/Crkl* were inactivated and found altered expression of many angiogenic genes and some signaling pathways such as NOTCH signaling, which help to explain the molecular mechanisms of observed phenotypes. Although peripheral vascular malformations have not been identified for Miller-Dieker syndrome as of yet, they have been identified in patients with 22q11.2DS.^94–96^ Our study provides insights into the cause of vasculature defects in humans.

## Acknowledgements

We thank Dr. Deyou Zheng from Albert Einstein College of Medicine for suggestions of selecting scRNA-seq sequencing strategy and identifying differentially expressed genes. We thank Dr. Tom Curran and Dr. Taeju Park from Children’s Research Institute (Children’s Mercy Kansas City) for the gift of *Crk* flox mouse strain and insightful comments for the manuscript. We very much appreciate input on our hematopoiesis experiments from Drs. Brian Black and Tanvi Sinha at the University of California, San Francisco. We thank Dr. Tao Wang from Albert Einstein College of Medicine for suggestions in choosing statistical methods. We thank Kevyn Jackson for her involvement in correcting error scripts when using Rstudio. We also want to thank the Flow Cytometry, Genomics, and Analytical Imaging core facilities at Albert Einstein College of Medicine.

## Sources of funding

Our research is supported by National Institutes of Health [P01 HD070454 (BEM, BZ); R01 HL132577 (BEM, BZ)]; R01 HL159515 (BZ, BEM), R01 HL157157 (BEM), U54 HD090260 (BEM), R01 HL157347 (BZ), and American Heart Association predoctoral fellowship [19PRE34380071 (LS)].

## Disclosures

None.

## Non-standard Abbreviations and Acronyms

EC: Endothelial cell
SMC: Smooth muscle cell
scRNA-seq: Single cell RNA sequencing
WHIF: Whole mount immunofluorescence
NICD: Notch1 intracellular domain
2-allele cKO embryo: Inactivate one allele of Crk and one allele of Crkl
3-allele cKO embryo: Inactivate one allele of Crk and two alleles of Crkl (and vice versa)
4-allele cKO embryo: Inactivate both alleles of Crk and Crkl
HPC: Hematopoietic progenitor cell
DEG: Differentially expressed gene
GO: Gene ontology

## Notes

### Competing Interest Statement

The authors have declared no competing interest.

## References

1. Feller SM. Crk family adaptors-signalling complex formation and biological roles. Oncogene. 2001;20:6348–6371. doi: 10.1038/sj.onc.1204779

2. Matsuda M, Tanaka S, Nagata S, Kojima A, Kurata T, Shibuya M. Two species of human CRK cDNA encode proteins with distinct biological activities. Mol Cell Biol. 1992;12:3482–3489. doi: 10.1128/mcb.12.8.3482-3489.1992

3. ten Hoeve J, Morris C, Heisterkamp N, Groffen J. Isolation and chromosomal localization of CRKL, a human crk-like gene. Oncogene. 1993;8:2469–2474.

4. Greenberg F, Courtney KB, Wessels RA, Huhta J, Carpenter RJ, Rich DC, Ledbetter DH. Prenatal diagnosis of deletion 17p13 associated with DiGeorge anomaly. Am J Med Genet. 1988;31:1–4. doi: 10.1002/ajmg.1320310102

5. Pilz DT, Quarrell OW. Syndromes with lissencephaly. J Med Genet. 1996;33:319–323. doi: 10.1136/jmg.33.4.319

6. Cardoso C, Leventer RJ, Ward HL, Toyo-Oka K, Chung J, Gross A, Martin CL, Allanson J, Pilz DT, Olney AH, et al. Refinement of a 400-kb critical region allows genotypic differentiation between isolated lissencephaly, Miller-Dieker syndrome, and other phenotypes secondary to deletions of 17p13.3. Am J Hum Genet. 2003;72:918–930. doi: 10.1086/374320

7. Edelmann L, Pandita RK, Morrow BE. Low-copy repeats mediate the common 3-Mb deletion in patients with velo-cardio-facial syndrome. Am J Hum Genet. 1999;64:1076–1086. doi: 10.1086/302343

8. Shaikh TH, Kurahashi H, Saitta SC, O’Hare AM, Hu P, Roe BA, Driscoll DA, McDonald-McGinn DM, Zackai EH, Budarf ML, et al. Chromosome 22-specific low copy repeats and the 22q11.2 deletion syndrome: genomic organization and deletion endpoint analysis. Hum Mol Genet. 2000;9:489–501. doi: 10.1093/hmg/9.4.489

9. Miller JQ. Lissencephaly in 2 Siblings. Neurology. 1963;13:841–850. doi: 10.1212/wnl.13.10.841

10. Saltzman DH, Krauss CM, Goldman JM, Benacerraf BR. Prenatal diagnosis of lissencephaly. Prenat Diagn. 1991;11:139–143. doi: 10.1002/pd.1970110302

11. Thomas MA, Duncan AM, Bardin C, Kaloustian VM. Lissencephaly with der(17)t(17;20)(p13.3;p12.2)mat. Am J Med Genet A. 2004;124A:292–295. doi: 10.1002/ajmg.a.20373

12. Nagamani SC, Zhang F, Shchelochkov OA, Bi W, Ou Z, Scaglia F, Probst FJ, Shinawi M, Eng C, Hunter JV, et al. Microdeletions including YWHAE in the Miller-Dieker syndrome region on chromosome 17p13.3 result in facial dysmorphisms, growth restriction, and cognitive impairment. J Med Genet. 2009;46:825–833. doi: 10.1136/jmg.2009.067637

13. Chen CP, Liu YP, Lin SP, Chen M, Tsai FJ, Chen YT, Chen LF, Hwang JK, Wang W. Ventriculomegaly, intrauterine growth restriction, and congenital heart defects as salient prenatal sonographic findings of Miller-Dieker lissencephaly syndrome associated with monosomy 17p (17p13.2 --> pter) in a fetus. Taiwan J Obstet Gynecol. 2010;49:81–86. doi: 10.1016/S1028-4559(10)60015-0

14. McDonald-McGinn DM, Kirschner R, Goldmuntz E, Sullivan K, Eicher P, Gerdes M, Moss E, Solot C, Wang P, Jacobs I, et al. The Philadelphia story: the 22q11.2 deletion: report on 250 patients. Genet Couns. 1999;10:11–24.

15. McDonald-McGinn DM, Sullivan KE, Marino B, Philip N, Swillen A, Vorstman JA, Zackai EH, Emanuel BS, Vermeesch JR, Morrow BE, et al. 22q11.2 deletion syndrome. Nat Rev Dis Primers. 2015;1:15071. doi: 10.1038/nrdp.2015.71

16. Imamoto A, Ki S, Li L, Iwamoto K, Maruthamuthu V, Devany J, Lu O, Kanazawa T, Zhang S, Yamada T, et al. Essential role of the Crk family-dosage in DiGeorge-like anomaly and metabolic homeostasis. Life Sci Alliance. 2020;3. doi: 10.26508/lsa.201900635

17. Racedo SE, McDonald-McGinn DM, Chung JH, Goldmuntz E, Zackai E, Emanuel BS, Zhou B, Funke B, Morrow BE. Mouse and human CRKL is dosage sensitive for cardiac outflow tract formation. Am J Hum Genet. 2015;96:235–244. doi: 10.1016/j.ajhg.2014.12.025

18. Park TJ, Boyd K, Curran T. Cardiovascular and craniofacial defects in Crk-null mice. Mol Cell Biol. 2006;26:6272–6282. doi: 10.1128/MCB.00472-06

19. Guris DL, Fantes J, Tara D, Druker BJ, Imamoto A. Mice lacking the homologue of the human 22q11.2 gene CRKL phenocopy neurocristopathies of DiGeorge syndrome. Nat Genet. 2001;27:293–298. doi: 10.1038/85855

20. Shi L, Racedo SE, Diacou A, Park T, Zhou B, Morrow BE. Crk and Crkl have shared functions in neural crest cells for cardiac outflow tract septation and vascular smooth muscle differentiation. Hum Mol Genet. 2022;31:1197–1215. doi: 10.1093/hmg/ddab313

21. Park T, Curran T. Requirement for Crk and CrkL during postnatal lens development. Biochem Biophys Res Commun. 2020;529:603–607. doi: 10.1016/j.bbrc.2020.06.108

22. Collins TN, Mao Y, Li H, Bouaziz M, Hong A, Feng GS, Wang F, Quilliam LA, Chen L, Park T, et al. Crk proteins transduce FGF signaling to promote lens fiber cell elongation. Elife. 2018;7. doi: 10.7554/eLife.32586

23. Park TJ, Curran T. Crk and Crk-like play essential overlapping roles downstream of disabled-1 in the Reelin pathway. J Neurosci. 2008;28:13551–13562. doi: 10.1523/JNEUROSCI.4323-08.2008

24. Saga Y, Miyagawa-Tomita S, Takagi A, Kitajima S, Miyazaki J, Inoue T. MesP1 is expressed in the heart precursor cells and required for the formation of a single heart tube. Development. 1999;126:3437–3447.

25. Saga Y, Kitajima S, Miyagawa-Tomita S. Mesp1 expression is the earliest sign of cardiovascular development. Trends Cardiovasc Med. 2000;10:345–352. doi: 10.1016/s1050-1738(01)00069-x

26. Donadon M, Santoro MM. The origin and mechanisms of smooth muscle cell development in vertebrates. Development. 2021;148. doi: 10.1242/dev.197384

27. Navaratnam V, Kaufman MH, Skepper JN, Barton S, Guttridge KM. Differentiation of the myocardial rudiment of mouse embryos: an ultrastructural study including freeze-fracture replication. J Anat. 1986;146:65–85.

28. Ji RP, Phoon CK, Aristizabal O, McGrath KE, Palis J, Turnbull DH. Onset of cardiac function during early mouse embryogenesis coincides with entry of primitive erythroblasts into the embryo proper. Circ Res. 2003;92:133–135. doi: 10.1161/01.res.0000056532.18710.c0

29. Udan RS, Culver JC, Dickinson ME. Understanding vascular development. Wiley Interdiscip Rev Dev Biol. 2013;2:327–346. doi: 10.1002/wdev.91

30. Drake CJ, Fleming PA. Vasculogenesis in the day 6.5 to 9.5 mouse embryo. Blood. 2000;95:1671–1679.

31. Blanco R, Gerhardt H. VEGF and Notch in tip and stalk cell selection. Cold Spring Harb Perspect Med. 2013;3:a006569. doi: 10.1101/cshperspect.a006569

32. Trindade A, Kumar SR, Scehnet JS, Lopes-da-Costa L, Becker J, Jiang W, Liu R, Gill PS, Duarte A. Overexpression of delta-like 4 induces arterialization and attenuates vessel formation in developing mouse embryos. Blood. 2008;112:1720–1729. doi: 10.1182/blood-2007-09-112748

33. Ridgway J, Zhang G, Wu Y, Stawicki S, Liang WC, Chanthery Y, Kowalski J, Watts RJ, Callahan C, Kasman I, et al. Inhibition of Dll4 signalling inhibits tumour growth by deregulating angiogenesis. Nature. 2006;444:1083–1087. doi: 10.1038/nature05313

34. Noguera-Troise I, Daly C, Papadopoulos NJ, Coetzee S, Boland P, Gale NW, Lin HC, Yancopoulos GD, Thurston G. Blockade of Dll4 inhibits tumour growth by promoting non-productive angiogenesis. Nature. 2006;444:1032–1037. doi: 10.1038/nature05355

35. Limbourg FP, Takeshita K, Radtke F, Bronson RT, Chin MT, Liao JK. Essential role of endothelial Notch1 in angiogenesis. Circulation. 2005;111:1826–1832. doi: 10.1161/01.CIR.0000160870.93058.DD

36. Gale NW, Dominguez MG, Noguera I, Pan L, Hughes V, Valenzuela DM, Murphy AJ, Adams NC, Lin HC, Holash J, et al. Haploinsufficiency of delta-like 4 ligand results in embryonic lethality due to major defects in arterial and vascular development. Proc Natl Acad Sci U S A. 2004;101:15949–15954. doi: 10.1073/pnas.0407290101

37. Fischer A, Schumacher N, Maier M, Sendtner M, Gessler M. The Notch target genes Hey1 and Hey2 are required for embryonic vascular development. Genes Dev. 2004;18:901–911. doi: 10.1101/gad.291004

38. Duarte A, Hirashima M, Benedito R, Trindade A, Diniz P, Bekman E, Costa L, Henrique D, Rossant J. Dosage-sensitive requirement for mouse Dll4 in artery development. Genes Dev. 2004;18:2474–2478. doi: 10.1101/gad.1239004

39. Lawson ND, Scheer N, Pham VN, Kim CH, Chitnis AB, Campos-Ortega JA, Weinstein BM. Notch signaling is required for arterial-venous differentiation during embryonic vascular development. Development. 2001;128:3675–3683. doi: 10.1242/dev.128.19.3675

40. Patel-Hett S, D’Amore PA. Signal transduction in vasculogenesis and developmental angiogenesis. Int J Dev Biol. 2011;55:353–363. doi: 10.1387/ijdb.103213sp

41. Saga Y, Hata N, Kobayashi S, Magnuson T, Seldin MF, Taketo MM. MesP1: a novel basic helix-loop-helix protein expressed in the nascent mesodermal cells during mouse gastrulation. Development. 1996;122:2769–2778.

42. Mao X, Fujiwara Y, Chapdelaine A, Yang H, Orkin SH. Activation of EGFP expression by Cre-mediated excision in a new ROSA26 reporter mouse strain. Blood. 2001;97:324–326.

43. Hao Y, Hao S, Andersen-Nissen E, Mauck WM, 3rd, Zheng S, Butler A, Lee MJ, Wilk AJ, Darby C, Zager M, et al. Integrated analysis of multimodal single-cell data. Cell. 2021;184:3573–3587 e3529. doi: 10.1016/j.cell.2021.04.048

44. Liu Y, Wang T, Zhou B, Zheng D. Robust integration of multiple single-cell RNA sequencing datasets using a single reference space. Nat Biotechnol. 2021;39:877–884. doi: 10.1038/s41587-021-00859-x

45. Chen J, Bardes EE, Aronow BJ, Jegga AG. ToppGene Suite for gene list enrichment analysis and candidate gene prioritization. Nucleic Acids Res. 2009;37:W305–311. doi: 10.1093/nar/gkp427

46. La Manno G, Soldatov R, Zeisel A, Braun E, Hochgerner H, Petukhov V, Lidschreiber K, Kastriti ME, Lonnerberg P, Furlan A, et al. RNA velocity of single cells. Nature. 2018;560:494–498. doi: 10.1038/s41586-018-0414-6

47. Bergen V, Lange M, Peidli S, Wolf FA, Theis FJ. Generalizing RNA velocity to transient cell states through dynamical modeling. Nat Biotechnol. 2020;38:1408–1414. doi: 10.1038/s41587-020-0591-3

48. Lange M, Bergen V, Klein M, Setty M, Reuter B, Bakhti M, Lickert H, Ansari M, Schniering J, Schiller HB, et al. CellRank for directed single-cell fate mapping. Nat Methods. 2022;19:159–170. doi: 10.1038/s41592-021-01346-6

49. De Val S, Black BL. Transcriptional control of endothelial cell development. Dev Cell. 2009;16:180–195. doi: 10.1016/j.devcel.2009.01.014

50. Lee D, Park C, Lee H, Lugus JJ, Kim SH, Arentson E, Chung YS, Gomez G, Kyba M, Lin S, et al. ER71 acts downstream of BMP, Notch, and Wnt signaling in blood and vessel progenitor specification. Cell Stem Cell. 2008;2:497–507. doi: 10.1016/j.stem.2008.03.008

51. Palis J, Robertson S, Kennedy M, Wall C, Keller G. Development of erythroid and myeloid progenitors in the yolk sac and embryo proper of the mouse. Development. 1999;126:5073–5084. doi: 10.1242/dev.126.22.5073

52. Yamane T. Mouse Yolk Sac Hematopoiesis. Front Cell Dev Biol. 2018;6:80. doi: 10.3389/fcell.2018.00080

53. Dzierzak E, Speck NA. Of lineage and legacy: the development of mammalian hematopoietic stem cells. Nat Immunol. 2008;9:129–136. doi: 10.1038/ni1560

54. Canu G, Ruhrberg C. First blood: the endothelial origins of hematopoietic progenitors. Angiogenesis. 2021;24:199–211. doi: 10.1007/s10456-021-09783-9

55. Zovein AC, Hofmann JJ, Lynch M, French WJ, Turlo KA, Yang Y, Becker MS, Zanetta L, Dejana E, Gasson JC, et al. Fate tracing reveals the endothelial origin of hematopoietic stem cells. Cell Stem Cell. 2008;3:625–636. doi: 10.1016/j.stem.2008.09.018

56. Swiers G, Rode C, Azzoni E, de Bruijn MF. A short history of hemogenic endothelium. Blood Cells Mol Dis. 2013;51:206–212. doi: 10.1016/j.bcmd.2013.09.005

57. Birge RB, Kalodimos C, Inagaki F, Tanaka S. Crk and CrkL adaptor proteins: networks for physiological and pathological signaling. Cell Commun Signal. 2009;7:13. doi: 10.1186/1478-811X-7-13

58. Hulsen T, de Vlieg J, Alkema W. BioVenn - a web application for the comparison and visualization of biological lists using area-proportional Venn diagrams. BMC Genomics. 2008;9:488. doi: 10.1186/1471-2164-9-488

59. Li JL, Sainson RC, Shi W, Leek R, Harrington LS, Preusser M, Biswas S, Turley H, Heikamp E, Hainfellner JA, et al. Delta-like 4 Notch ligand regulates tumor angiogenesis, improves tumor vascular function, and promotes tumor growth in vivo. Cancer Res. 2007;67:11244–11253. doi: 10.1158/0008-5472.CAN-07-0969

60. Uyttendaele H, Ho J, Rossant J, Kitajewski J. Vascular patterning defects associated with expression of activated Notch4 in embryonic endothelium. Proc Natl Acad Sci U S A. 2001;98:5643–5648. doi: 10.1073/pnas.091584598

61. Uyttendaele H, Marazzi G, Wu G, Yan Q, Sassoon D, Kitajewski J. Notch4/int-3, a mammary proto-oncogene, is an endothelial cell-specific mammalian Notch gene. Development. 1996;122:2251–2259. doi: 10.1242/dev.122.7.2251

62. Li W, Liu C, Burns N, Hayashi J, Yoshida A, Sajja A, Gonzalez-Hernandez S, Gao JL, Murphy PM, Kubota Y, et al. Alterations in the spatiotemporal expression of the chemokine receptor CXCR4 in endothelial cells cause failure of hierarchical vascular branching. Dev Biol. 2021;477:70–84. doi: 10.1016/j.ydbio.2021.05.008

63. Ma Q, Jones D, Borghesani PR, Segal RA, Nagasawa T, Kishimoto T, Bronson RT, Springer TA. Impaired B-lymphopoiesis, myelopoiesis, and derailed cerebellar neuron migration in CXCR4- and SDF-1-deficient mice. Proc Natl Acad Sci U S A. 1998;95:9448–9453. doi: 10.1073/pnas.95.16.9448

64. Tachibana K, Hirota S, Iizasa H, Yoshida H, Kawabata K, Kataoka Y, Kitamura Y, Matsushima K, Yoshida N, Nishikawa S, et al. The chemokine receptor CXCR4 is essential for vascularization of the gastrointestinal tract. Nature. 1998;393:591–594. doi: 10.1038/31261

65. John H, Radtke K, Standker L, Forssmann WG. Identification and characterization of novel endogenous proteolytic forms of the human angiogenesis inhibitors restin and endostatin. Biochim Biophys Acta. 2005;1747:161–170. doi: 10.1016/j.bbapap.2004.10.013

66. Ramchandran R, Dhanabal M, Volk R, Waterman MJ, Segal M, Lu H, Knebelmann B, Sukhatme VP. Antiangiogenic activity of restin, NC10 domain of human collagen XV: comparison to endostatin. Biochem Biophys Res Commun. 1999;255:735–739. doi: 10.1006/bbrc.1999.0248

67. Takeda N, Maemura K, Imai Y, Harada T, Kawanami D, Nojiri T, Manabe I, Nagai R. Endothelial PAS domain protein 1 gene promotes angiogenesis through the transactivation of both vascular endothelial growth factor and its receptor, Flt-1. Circ Res. 2004;95:146–153. doi: 10.1161/01.RES.0000134920.10128.b4

68. Peng J, Zhang L, Drysdale L, Fong GH. The transcription factor EPAS-1/hypoxia-inducible factor 2alpha plays an important role in vascular remodeling. Proc Natl Acad Sci U S A. 2000;97:8386–8391. doi: 10.1073/pnas.140087397

69. Kobayashi S, Yamashita T, Ohneda K, Nagano M, Kimura K, Nakai H, Poellinger L, Ohneda O. Hypoxia-inducible factor-3alpha promotes angiogenic activity of pulmonary endothelial cells by repressing the expression of the VE-cadherin gene. Genes Cells. 2015;20:224–241. doi: 10.1111/gtc.12215

70. Brugarolas J, Lei K, Hurley RL, Manning BD, Reiling JH, Hafen E, Witters LA, Ellisen LW, Kaelin WG, Jr. Regulation of mTOR function in response to hypoxia by REDD1 and the TSC1/TSC2 tumor suppressor complex. Genes Dev. 2004;18:2893–2904. doi: 10.1101/gad.1256804

71. Leveen P, Pekny M, Gebre-Medhin S, Swolin B, Larsson E, Betsholtz C. Mice deficient for PDGF B show renal, cardiovascular, and hematological abnormalities. Genes Dev. 1994;8:1875–1887. doi: 10.1101/gad.8.16.1875

72. Caron KM, Smithies O. Extreme hydrops fetalis and cardiovascular abnormalities in mice lacking a functional Adrenomedullin gene. Proc Natl Acad Sci U S A. 2001;98:615–619. doi: 10.1073/pnas.98.2.615

73. Shindo T, Kurihara Y, Nishimatsu H, Moriyama N, Kakoki M, Wang Y, Imai Y, Ebihara A, Kuwaki T, Ju KH, et al. Vascular abnormalities and elevated blood pressure in mice lacking adrenomedullin gene. Circulation. 2001;104:1964–1971. doi: 10.1161/hc4101.097111

74. Shimosawa T, Shibagaki Y, Ishibashi K, Kitamura K, Kangawa K, Kato S, Ando K, Fujita T. Adrenomedullin, an endogenous peptide, counteracts cardiovascular damage. Circulation. 2002;105:106–111. doi: 10.1161/hc0102.101399

75. Wetzel-Strong SE, Li M, Klein KR, Nishikimi T, Caron KM. Epicardial-derived adrenomedullin drives cardiac hyperplasia during embryogenesis. Dev Dyn. 2014;243:243–256. doi: 10.1002/dvdy.24065

76. De Val S, Chi NC, Meadows SM, Minovitsky S, Anderson JP, Harris IS, Ehlers ML, Agarwal P, Visel A, Xu SM, et al. Combinatorial regulation of endothelial gene expression by ets and forkhead transcription factors. Cell. 2008;135:1053–1064. doi: 10.1016/j.cell.2008.10.049

77. Oh SY, Kim JY, Park C. The ETS Factor, ETV2: a Master Regulator for Vascular Endothelial Cell Development. Mol Cells. 2015;38:1029–1036. doi: 10.14348/molcells.2015.0331

78. del Toro R, Prahst C, Mathivet T, Siegfried G, Kaminker JS, Larrivee B, Breant C, Duarte A, Takakura N, Fukamizu A, et al. Identification and functional analysis of endothelial tip cell-enriched genes. Blood. 2010;116:4025–4033. doi: 10.1182/blood-2010-02-270819

79. Helker CS, Eberlein J, Wilhelm K, Sugino T, Malchow J, Schuermann A, Baumeister S, Kwon HB, Maischein HM, Potente M, et al. Apelin signaling drives vascular endothelial cells toward a pro-angiogenic state. Elife. 2020;9. doi: 10.7554/eLife.55589

80. Shutter JR, Scully S, Fan W, Richards WG, Kitajewski J, Deblandre GA, Kintner CR, Stark KL. Dll4, a novel Notch ligand expressed in arterial endothelium. Genes Dev. 2000;14:1313–1318.

81. Risau W, Flamme I. Vasculogenesis. Annu Rev Cell Dev Biol. 1995;11:73–91. doi: 10.1146/annurev.cb.11.110195.000445

82. Salameh A, Galvagni F, Bardelli M, Bussolino F, Oliviero S. Direct recruitment of CRK and GRB2 to VEGFR-3 induces proliferation, migration, and survival of endothelial cells through the activation of ERK, AKT, and JNK pathways. Blood. 2005;106:3423–3431. doi: 10.1182/blood-2005-04-1388

83. Stoletov KV, Gong C, Terman BI. Nck and Crk mediate distinct VEGF-induced signaling pathways that serve overlapping functions in focal adhesion turnover and integrin activation. Exp Cell Res. 2004;295:258–268. doi: 10.1016/j.yexcr.2004.01.008

84. Shawber CJ, Kitajewski J. Notch function in the vasculature: insights from zebrafish, mouse and man. Bioessays. 2004;26:225–234. doi: 10.1002/bies.20004

85. Krebs LT, Xue Y, Norton CR, Shutter JR, Maguire M, Sundberg JP, Gallahan D, Closson V, Kitajewski J, Callahan R, et al. Notch signaling is essential for vascular morphogenesis in mice. Genes Dev. 2000;14:1343–1352.

86. Xue Y, Gao X, Lindsell CE, Norton CR, Chang B, Hicks C, Gendron-Maguire M, Rand EB, Weinmaster G, Gridley T. Embryonic lethality and vascular defects in mice lacking the Notch ligand Jagged1. Hum Mol Genet. 1999;8:723–730. doi: 10.1093/hmg/8.5.723

87. Krebs LT, Shutter JR, Tanigaki K, Honjo T, Stark KL, Gridley T. Haploinsufficient lethality and formation of arteriovenous malformations in Notch pathway mutants. Genes Dev. 2004;18:2469–2473. doi: 10.1101/gad.1239204

88. Park T, Koptyra M, Curran T. Fibroblast Growth Requires CT10 Regulator of Kinase (Crk) and Crk-like (CrkL). The Journal of biological chemistry. 2016;291:26273–26290. doi: 10.1074/jbc.M116.764613

89. Park TJ, Curran T. Essential roles of Crk and CrkL in fibroblast structure and motility. Oncogene. 2014;33:5121–5132. doi: 10.1038/onc.2013.453

90. Cohen M, Georgiou M, Stevenson NL, Miodownik M, Baum B. Dynamic filopodia transmit intermittent Delta-Notch signaling to drive pattern refinement during lateral inhibition. Dev Cell. 2010;19:78–89. doi: 10.1016/j.devcel.2010.06.006

91. De Joussineau C, Soule J, Martin M, Anguille C, Montcourrier P, Alexandre D. Delta-promoted filopodia mediate long-range lateral inhibition in Drosophila. Nature. 2003;426:555–559. doi: 10.1038/nature02157

92. Gomez-Lamarca MJ, Cobreros-Reguera L, Ibanez-Jimenez B, Palacios IM, Martin-Bermudo MD. Integrins regulate epithelial cell differentiation by modulating Notch activity. J Cell Sci. 2014;127:4667–4678. doi: 10.1242/jcs.153122

93. Rallis C, Pinchin SM, Ish-Horowicz D. Cell-autonomous integrin control of Wnt and Notch signalling during somitogenesis. Development. 2010;137:3591–3601. doi: 10.1242/dev.050070

94. Gokturk B, Topcu-Yilmaz P, Bozkurt B, Yildirim MS, Guner SN, Sayar EH, Reisli I. Ocular Findings in Children With 22q11.2 Deletion Syndrome. J Pediatr Ophthalmol Strabismus. 2016;53:218–222. doi: 10.3928/01913913-20160427-01

95. Forbes BJ, Binenbaum G, Edmond JC, DeLarato N, McDonald-McGinn DM, Zackai EH. Ocular findings in the chromosome 22q11.2 deletion syndrome. J AAPOS. 2007;11:179–182. doi: 10.1016/j.jaapos.2006.08.006

96. Hill VE, Pietucha S, Ells AL. Congenital vascular tortuosity in DiGeorge syndrome mimicking significant retinopathy of prematurity. Arch Ophthalmol. 2004;122:132–133. doi: 10.1001/archopht.122.1.132

